# The plasma degradome reflects later development of NASH fibrosis after liver transplant

**DOI:** 10.1101/2023.01.30.526241

**Authors:** Jiang Li, Toshifumi Sato, María Hernández-Tejero, Juliane I. Beier, Khaled Sayed, Panayiotis V Benos, Daniel W Wilkey, Abhinav Humar, Michael L Merchant, Andres Duarte-Rojo, Gavin E Arteel

**Affiliations:** Department of Medicine, University of Pittsburgh, Pittsburgh, PA, USA; Pittsburgh Liver Research Center, University of Pittsburgh, Pittsburgh, PA, USA; Department of Epidemiology, University of Florida, Gainesville, FL, USA; Department of Medicine, University of Louisville, Louisville, KY, USA; Department of Surgery, University of Pittsburgh, Pittsburgh, PA, USA; Division of Gastroenterology and Hepatology, Northwestern University, Chicago, IL; Comprehensive Transplant Center, Northwestern Medicine and Feinberg School of Medicine, Northwestern University, Chicago, IL

**Author notes:** Send all correspondence to: Gavin E. Arteel, PhD, FAASLD, Thomas E. Starzl Biomedical Science Tower, West 1143, 200 Lothrop Street, Pittsburgh, PA 15213.

**Keywords:** Extracellular matrix, remodeling, liver transplantation outcomes, translational study

## Abstract

Although liver transplantation (LT) is an effective therapy for cirrhosis, the risk of post-LT NASH is alarmingly high and is associated with accelerated progression to fibrosis/cirrhosis, cardiovascular disease, and decreased survival. Lack of risk stratification strategies hamper liver undergoes significant remodeling during inflammatory injury. During such remodeling, degraded peptide fragments (i.e., ‘degradome’) of the ECM and other proteins increase in plasma, making it a useful diagnostic/prognostic tool in chronic liver disease. To investigate whether inflammatory liver injury caused by post-LT NASH would yield a unique degradome profile, predictive of severe post-LT NASH fibrosis, we performed a retrospective analysis of 22 biobanked samples from the Starzl Transplantation Institute (12 with post-LT NASH after 5 years and 10 without). Total plasma peptides were isolated and analyzed by 1D-LC-MS/MS analysis using a Proxeon EASY-nLC 1000 UHPLC and nanoelectrospray ionization into an Orbitrap Elite mass spectrometer. Qualitative and quantitative peptide features data were developed from MSn datasets using PEAKS Studio X (v10). LC-MS/MS yielded ∼2700 identifiable peptide features based on the results from Peaks Studio analysis. Several peptides were significantly altered in patients that later developed fibrosis and heatmap analysis of the top 25 most significantly-changed peptides, most of which were ECM-derived, clustered the 2 patient groups well. Supervised modeling of the dataset indicated that a fraction of the total peptide signal (∼15%) could explain the differences between the groups, indicating a strong potential for representative biomarker selection. A similar degradome profile was observed when the plasma degradome patterns were compared being obesity sensitive (C57Bl6/J) and insensitive (AJ) mouse strains. Both The plasma degradome profile of post-LT patients yields stark difference based on later development of post-LT NASH fibrosis. This approach could yield new “fingerprints” that can serve as minimally-invasive biomarkers of negative outcomes post-LT.

## Introduction

Liver transplantation (LT) is an increasingly utilized therapeutic approach for advanced liver disease, regardless of etiology. Cirrhosis from non-alcoholic steatohepatitis (NASH) is now one of the most common indications for liver transplantation (LT) in the US.^1,2^ Although LT is an effective therapy for NASH, the risk of post-transplant recurrence is alarmingly high;^3,4^ recurrent NASH has an incidence of up to 70% at 5 years, when per protocol biopsies are performed.^5^ While the presence of simple steatosis does not appear to impact overall graft and patient survival,^3^ the presence of NASH is associated with accelerated progression to fibrosis/cirrhosis and cardiovascular disease (CVD), which negatively impact survival.^6,7^ Patients that develop de novo NASH, although they have a lower incidence (17% at 5 years), are also at high risk for accelerated progression to NASH fibrosis.^5^ Indeed, 8% of per protocol biopsies already have post-LT NASH 1 year after living donor LT in our Center, irrespective of the original cause of liver disease, suggesting that the processes leading to post-LT NASH occur early in the post-LT period (Humar and colleagues, unpublished observations). Although lifestyle and drug therapies are promising approaches to decrease fibrosis related to NASH post-LT and its severe health effects, lack of effective strategies to stratify risk hamper the use of these tools. Moreover, post-LT NASH patients have been excluded from clinical trials testing novel drug therapies and as such, timely risk stratification followed by lifestyle interventions and management of comorbidities will remain as the standard of care in the foreseeable future.

The extracellular matrix (ECM) contains a diverse range of components that work bi-directionally with surrounding cells to create a dynamic and responsive microenvironment that regulates cell and tissue homeostasis.^8^ Proteases and protease inhibitors contribute to maintaining ECM homeostasis, as well as mediating its changes in response to stress or injury.^9,10^ Hepatic fibrosis is a canonical example of ECM dyshomeostasis, leading to accumulation of fibrillary ECM, such as collagen. Although hepatic fibrosis is considered almost synonymous to collagen accumulation,^11^ the qualitative and quantitative alterations to the hepatic ECM during fibrosis are much more diverse and certainly occur much earlier than fibrotic scarring of the liver.^12^ Indeed, transitional changes to the hepatic ECM appear to be key to the normal response to injury.^8^ Homeostasis in the ECM is mediated by a balance in the production of ECM, as well as in the degradation of existing ECM by matrix metalloproteinases (MMPs).^13^ Even in cases where there is a net increase in ECM in the liver (e.g., fibrosis) overall turnover is also increased.^14^ We and others have demonstrated that even acute liver injury causes robust changes to the hepatic ECM.

The peptidome is defined as the population of low molecular weight biologic peptides (e.g., 0.5-3 kDa), within the cells and biologic fluids. These compartments contain peptides critical for normal organismal function (e.g., peptide hormones and peptide neurotransmitters). The study of the peptidome (i.e., ‘peptidomics’) is a discipline related to proteomics, but with significant methodological and analytical differences.^15^ For example, as the original structure of the peptides is of interest, samples are not trypsin digested prior to mass spectrometry (MS) analysis, as is the case for bottom-up proteomic approaches. This difference originally limited peptide identification, as the available sequence assignment processes were based on trypsin-digested peptide databases; however, the development of de novo sequence assignment methods has addressed this concern.^16^ In addition to these small peptides, the peptidome also contains fragments of larger proteins degraded by normal and/or abnormal processes (i.e., ‘degradome’). The latter subset of the peptidome has generated great interest in some areas of human health as potential (surrogate) biomarkers for disease; for example, in cancer metastasis, and by extension, in overall patient outcome.^16,17^

The purpose of the current study was to identify plasma degradome profile differences in patients that developed post-LT NASH fibrosis to those that did not at an early time-point prior to detectable liver disease. The results in the human study were also compared to a preclinical model of NAFLD/NASH employing sensitive and insensitive strains of mice.

## Results

### Characterization of the degradome in transplant recipients

Patients were followed for a median of 2 years (range: 1 to 9 years). Twenty out of 22 (91%) cases had at least one liver biopsy available for review. Post-LT NAFLD was observed among 9 (41%) recipients whereas NASH developed in 6 (27%); 4 were classified as de novo NAFLD/NASH. Progression to at least significant fibrosis (F≥2 progressors) occurred in 12 recipients, 4 of whom had underlying post-LT NASH (3 de novo) and one with NAFLD based on a proton-density fat fraction of 29%. When comparing F≥2 progressors to non-progressors, there were no differences in key baseline characteristics. Importantly, there were no patients with diabetes mellitus at baseline. Among the group of progressors, 4 recipients (33%) ultimately developed cirrhosis. Allograft rejection, either T-cell- or antibody-mediated occurred on 3 cases (including 2 progressors with hepatitis C at baseline), none of whom had post-LT NAFLD or NASH.

### Analysis of LC-MS/MS data

After LC-MS/MS analysis (see Methods), Peaks was used for visualization of the MS data. Across the two patient groups, 2694 peptides were identified, corresponding to 721 distinct parent proteins (see supplemental Table 3). Of these, a total of 542 peptides (422 in fibrotic and 120 in nonfibrotic), corresponding to 179 distinct proteins, were changed at least 2-fold between patients who progressed onto ≥F2 fibrosis (“progressors”) versus those that did not (“non-progressors”). These changes are summarized graphically as a volcano plot (Figure 1A). Heatmap analysis and hierarchical clustering of top 25 changed peptides of individual patients (Figure 1B) clustered the “progressors” against the “non-progressors” well. Panel C. Supervised orthogonal partial least squares-discriminant analysis (OPLS-DA) score plot of the plasma peptidome profiles of individual patients (Figure 1C) indicated that a fraction (∼15%) of the total peptide could explain the differences between the groups.

**Figure 1.**
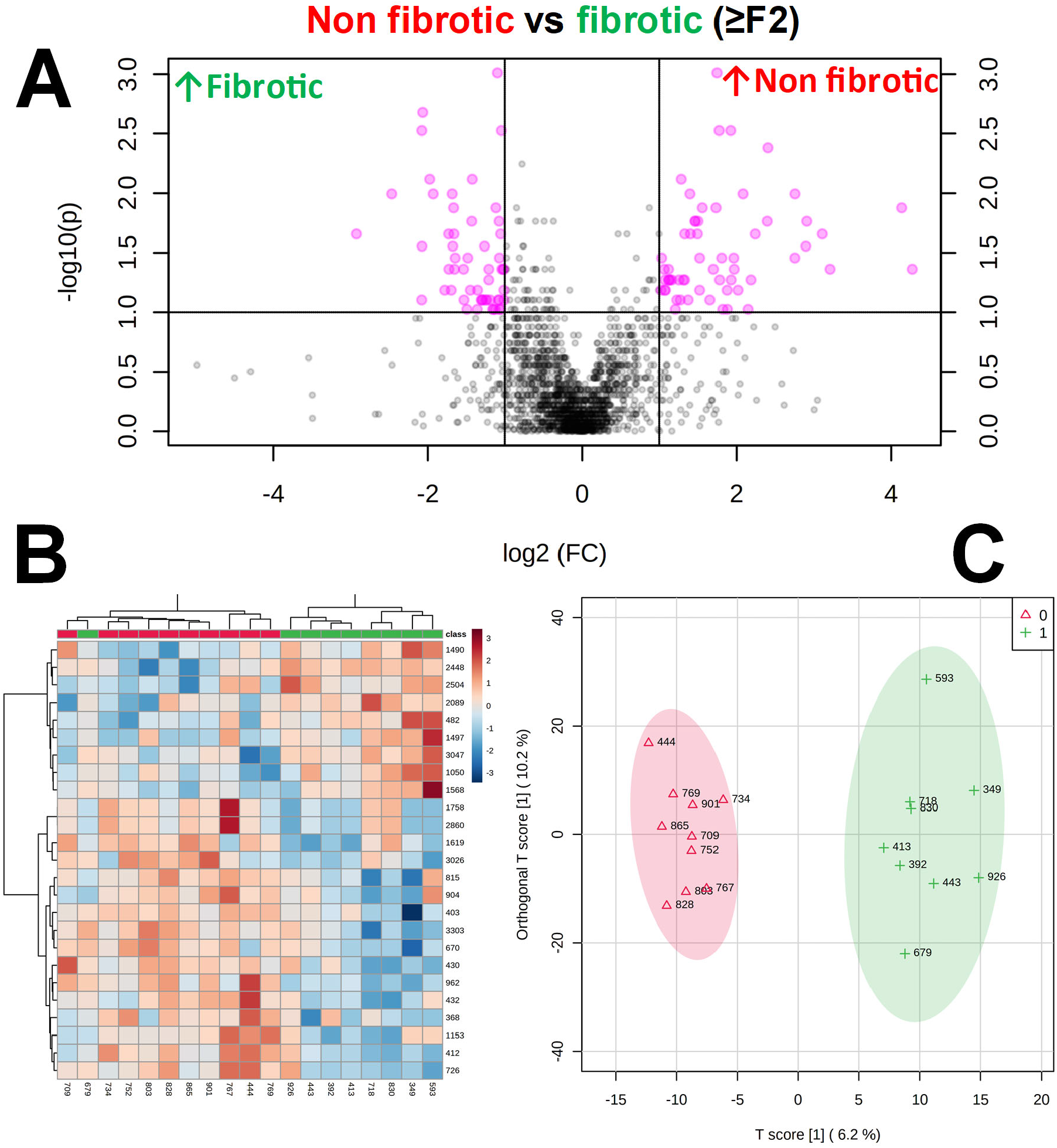
The plasma degradome in post-LT NASH fibrosis. Analysis of the peptidome/degradome in post-LT NASH fibrosis. Summary of analysis of fibrotic (≥F2; n=9; green) versus non-fibrotic (n=10; red) livers of post-LT patients 1 year after LT. **Panel A**. Volcano plots of changed peptides. **Panel B**. Heatmap and dendrogram of top 25 changed peptides of individual patients. **Panel C**. Orthogonal partial least squares-discriminant analysis (OPLS-DA) score plot of the plasma peptidome profiles of individual patients.

The parent proteins that contributed to the pattern of significantly changed peptides were analyzed by StringDB.^18,19^ This database queries physical and functional protein-protein associations that integrates both experimental and predicted associations. The output creates both graphical and categorical enrichment patterns in the queried dataset. Four key protein family clusters were enriched in the plasma degradome of patients that progressed to at least significant fibrosis (i.e., ≥F2 staging) (Figure 2A), including mitochondrial respiratory chain complex 1 (p=1.1×10^−10^), tubulin (p=7.4×10^−9^) collagen chains (p=7.7×10^−7^) and platelet α-granule (p=1.4×10^−6^). Similar pathways related to altered metabolism and remodeling were also enriched also in GO (Molecular function) and KEGG analyses (Figure 2B). GO terms for Cellular Component and Biological Processes provided by StringDB showed similar patterns (supplemental Tables 4 and 5, respectively).

**Figure 2.**
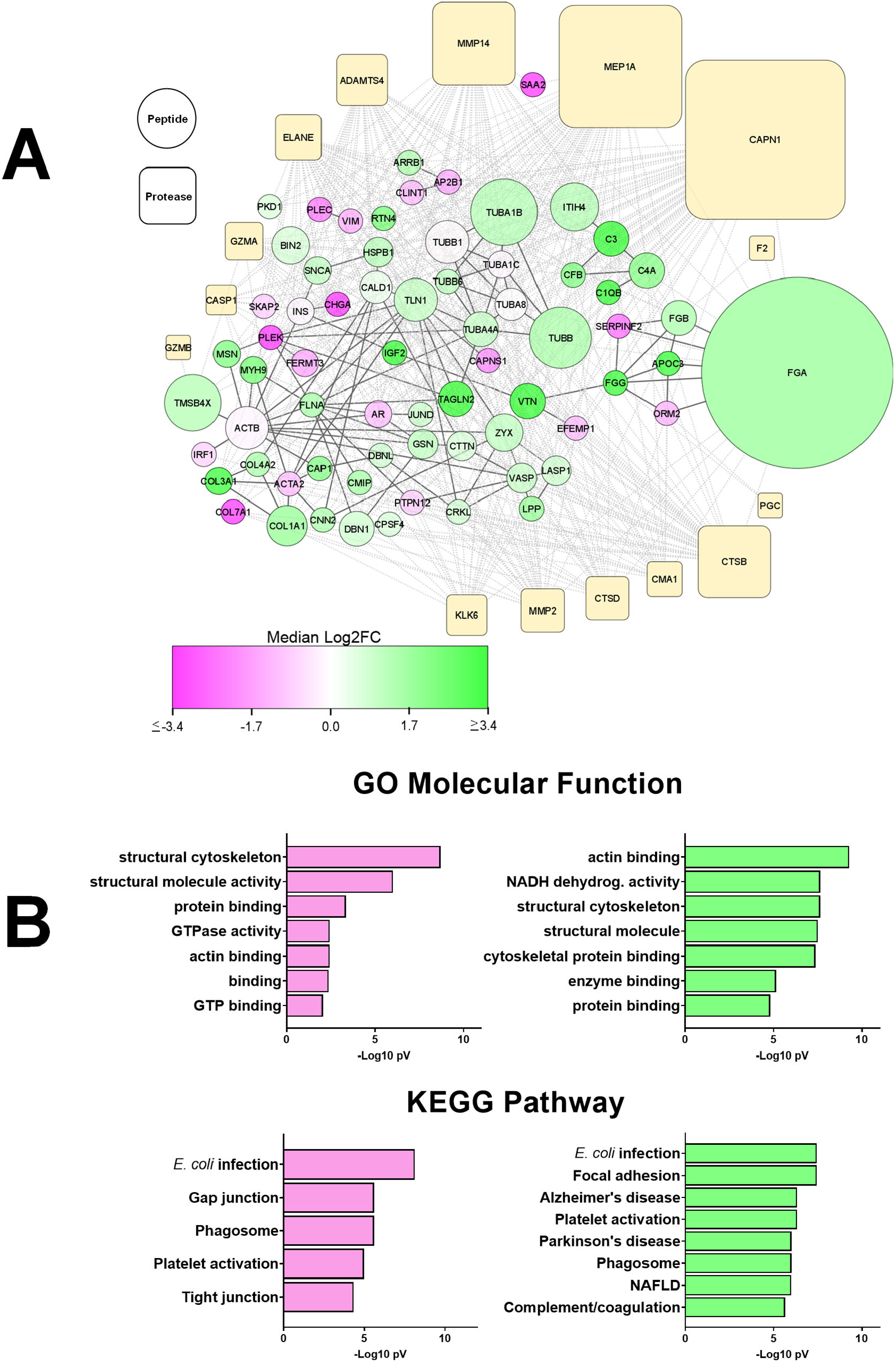
Cluster analysis of the peptidome/degradome in post-LT NASH fibrosis. The peptides significantly increased in NASH fibrosis (see Figure 1) were analyzed by the Proteasix (http://www.proteasix.org) algorithm using with the positive predictive value (PPV) cut-off to 80%. Protein-protein interaction network analysis of regulated proteomic data sets (q-value <0.05) was performed using Search Tool for the Retrieval of Interacting Genes/Proteins, STRING v11,^19^ with the highest confidence score (0.900). The resultant matrix of both Proteasix and STRING analysis were visualized using Cytoscape v3.9.1 (**Panel A**). Node sizes of the predicted proteases represented the relative frequency with which the top 15 proteases were predicted to mediate the observed cleavage (0.2-25%). Node sizes of the peptides represented the relative number of unique peptides (1-56) identified from each parent protein. Node color of the peptides represented the median Log2FC vs non-fibrosers for all peptides derived from that parent protein. Solid lines depict connections between the parent proteins identified by STRING; broken lines depict predicted protease events identified by Proteasix. (**Panel B**) Top GO (molecular functions) and KEGG terms identified cluster/pathway results of STRING analysis for decreased (pink) and increased (green) peptides.

## Comparing experimental NASH in sensitive and insensitive mice

Although other factors (e.g., immunosuppressive regimens) may also contribute, the primary risk factors for post-LT NASH with fibrosis appear to be synonymous with pre-LT NASH (e.g., obesity, metabolic syndrome and insulin resistance).^20-22^ Although no mouse model completely recapitulates human NAFLD/NASH, dietary manipulation with the obesogenic “Western” diet generates a phenotype similar to that found in human disease in sensitive mouse strains and have been used as tools to compare low-versus high-risk human populations.^23,24^ Accordingly, C57Bl6/J and AJ mice were fed a high-fat “Western” diet (HFD) or low-fat control diet (LFD) for 12 weeks. As expected, whereas C57Bl6/J mice were sensitive to the obesogenic effects of HFD, AJ mice were relatively insensitive. Specifically, C57Bl6/J mice gained >70% of their initial body weight (BW) when fed HFD, vs ∼20% for LFD (Figure 3A; supplemental Table 6). In contrast, HFD did not significantly alter the rate of BW gain in AJ mice, which was ∼30% over the course of the study (Figure 3A). Moreover whereas, HFD increased the homeostatic model assessment for insulin resistance (i.e., HOMA-IR) in C57Bl6/J mice, it did not in AJ mice (Figure 3A).

**Figure 3.**
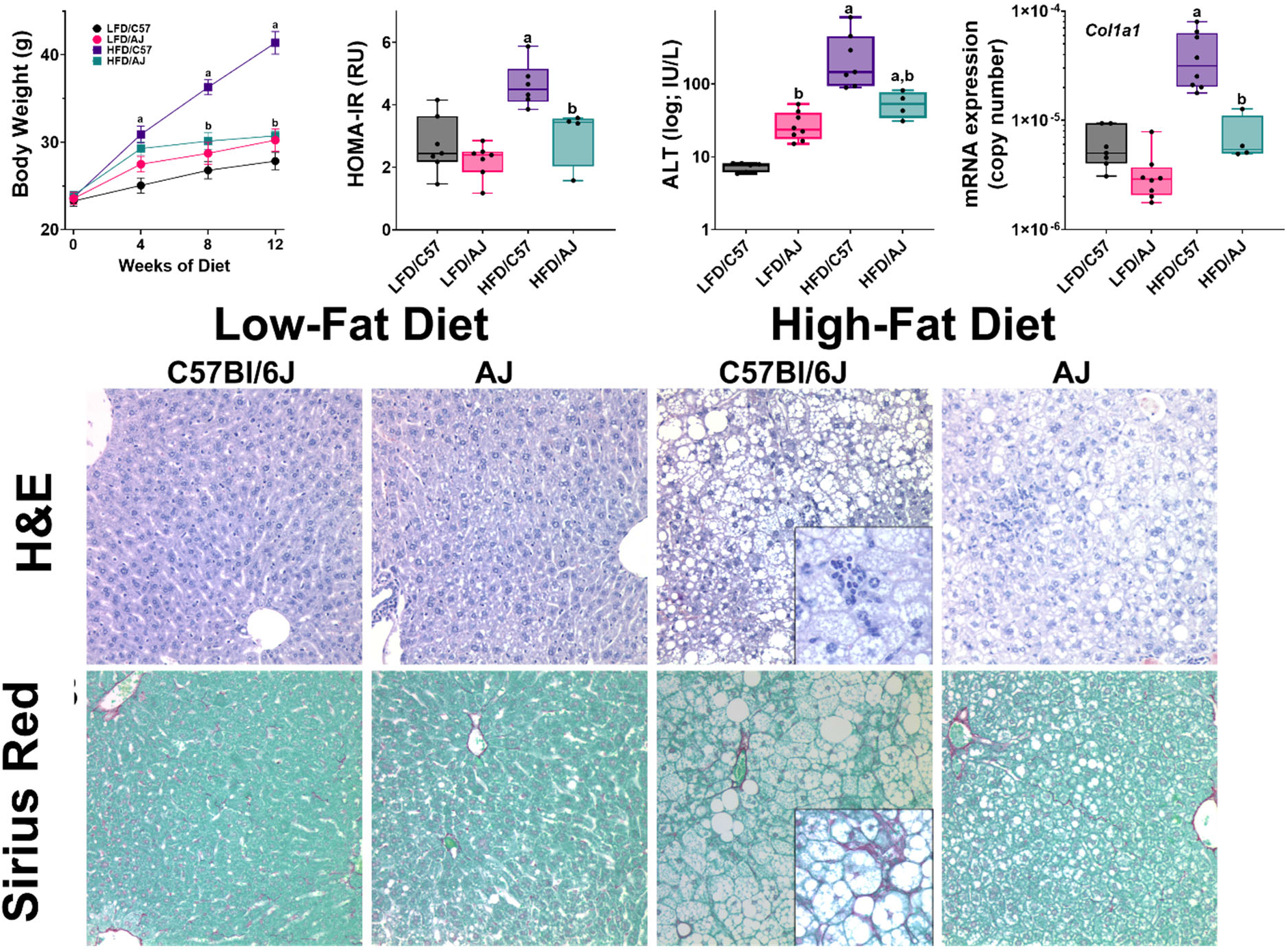
Strain differences in sensitivity to experimental NAFLD/NASH. C57Bl6/J (“C57”) and A/J (“AJ”) mice were fed the high-fat ‘Western’ (“HFD”) or low-fat control (“LFD”) diet for 12 weeks. The effect of HFD on BW, HOMA-IR plasma ALT (log scale) and expression of Col1A1 (log copy number) was compared (**Panel A**). **Panel B** shows representative photomicrographs of H&E (upper) and Sirius Red (lower) stains. Summary data (**Panel A**) are box and whisker plots (n=4-8/group), which show the median (thick line), interquartile range (box) and range (whiskers). ^a^*P <*0.05 for effect of HFD, ^b^P <0.05 for effect of mouse strain by 2-way ANOVA using Tukey’s post hoc test.

Sensitive C57Bl6/J mice also developed significant liver injury when fed a HFD, whereas AJ mice did not. HFD increased liver weight in C57Bl6/J mice by 85%, whereas liver weight did not change in AJ (Supplemental Table 6). HFD also caused severe steatosis, as determined by histological assessment (Figure 3B) and by biochemical analysis of lipid accumulation (Supplemental Table 6); although moderate macrovesicular steatosis and lipid accumulation was also observed in AJ mice fed HFD (Figure 3B, Supplemental Table 6), this effect was much less pronounced. The increase in hepatic transaminases caused by HFD (Figure 3A) was also more severe in C57Bl6/J compared to AJ mice. HFD induced expression of indices of fibrogenesis and collagen accumulation in C57Bl6/J mice (Figure 3A and Supplemental Figure 1) and caused the formation of “chicken-wire” fibrosis within the hepatic lobule (Figure 3B, inset).

Analogous to what we observed in post-LT patients (Figure 1), HFD dramatically altered the degradome profile in NASH sensitive C57 mice, as compared to the insensitive AJ mice (Figure 4A). Although unsupervised Principle Component Analysis (PCA) clustered both HFD- and LFD-fed AJ mice together, HFD and LFD feeding clustered the degradome pattern separately for C57BI6/J mice (figure 4A). Interestingly, heatmap (Figure 4C) and OPLS-DA (Figure 4D) analysis indicated several overlapping ‘hits’ (e.g., degraded tubulin, fibrinogen and collagen proteins) as found in the post-LT patients that developed ≥F2 NASH fibrosis (Figure 2). Indeed, as in the human data, fibrinogen (FIBA) protein products were dominant in the top scored peptides (Figure 4C).

**Figure 4.**
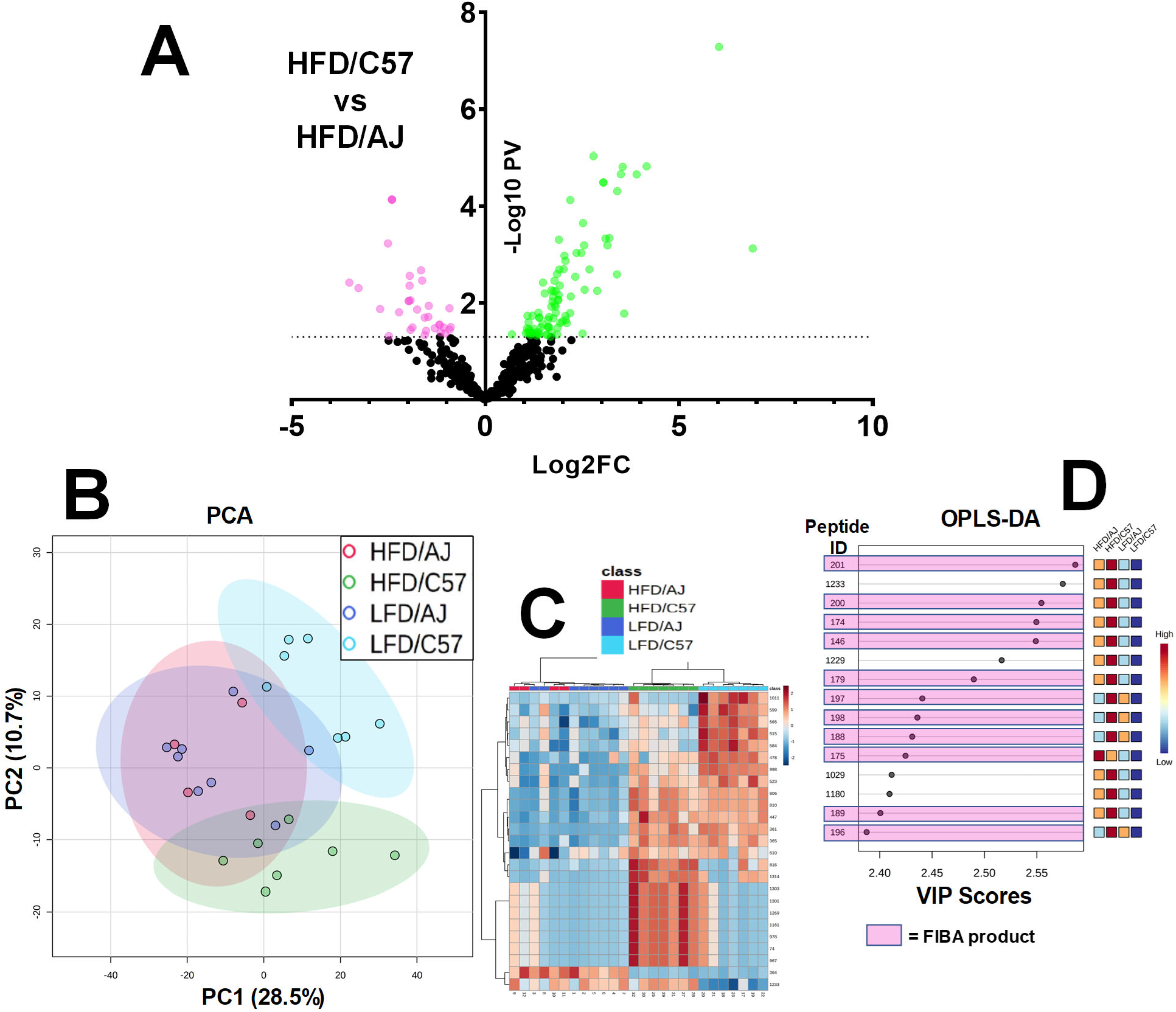
Mouse strain differences in the degradome profile in response to HFD. Feeding conditions are as described in Figure 4. **Panel A** shows a volcano plot comparing the plasma degradome profile between C57 to AJ mice fed HFD for 12 weeks. **Panel B** shows Principal Component Analysis (PCA) of the degradome profile in AJ and C57 mice fed low- or high-fat diets (LFD and HFD). **Panel C** shows, heatmap with dendrogram of the top 35 altered peptides. **Panel D** shows Variable Importance in Projection (VIP) scores of the top 15 peptides by supervised discriminant analysis (OPLS-DA; see also Figure 1). Pink highlighted peptides are derived from fibrinogen A (FBA).

Many proteases cleave substrates with high specificity only at certain sequence sites. Thus, information on the fragment sequence of degraded proteins can inform on proteases that may have generated this pattern. Proteasix (proteasix.org), an open-source peptide-centric tool to predict in silico the proteases involved in generating these peptides.^25^ Figure 2A also shows the relative frequency with which the top 15 proteases were predicted to generate the resultant degradome peptides by this analysis. Figure 5A shows a frequency distribution figure comparing the degradome in humans that progressed to ≥F2 after LT versus sensitive mice (C57Bl6/J) fed a high-fat diet. The two top predicted proteases, Calpain -1/-2 (Capn1/2) and Meprin-1a (Mep1a) were shared between the species under these conditions (Figure 2A and Figure 5A). The expression of the splice variants of Capn1/2 (*Capn1* and *Capn1*) was determined via real-time rtPCR in mouse liver (Figure 5B). Both isoform variants were expressed in liver, with *Capn2* expression being more abundant. HFD feeding significantly induced expression of these splice variants in the C57Bl6/J strain (Figure 5B). A similar effect was observed for *Mep1a* expression (Figure 5B). Several of the predicted CAPN1/2 and MEP1A substrates for these enzymes (Figure 5C and 5E) were analogous to the key nodes identified by StringDB analysis for the degradome as a whole (Figure 2A) and the enriched GO Biological Processes of these substrates related to remodeling and coagulation cascade activation (Figure 5D).

**Figure 5.**
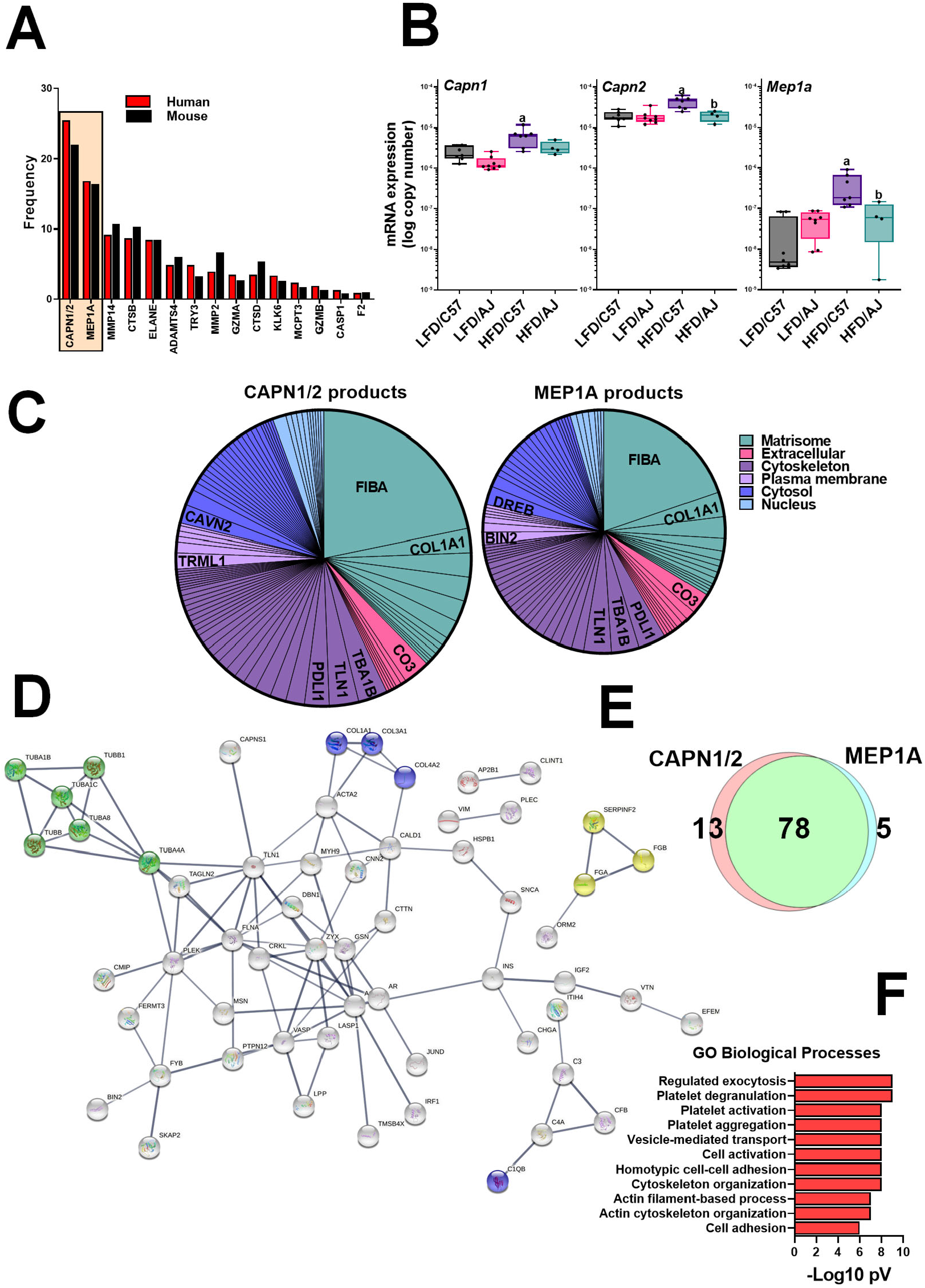
The plasma degradome predicts hepatic protease induction in both human pre-LT NASH fibrosis and experimental mouse NAFLD/NASH. **Panel A** shows the top 15 proteases by predicted Proteasix analysis in post-LT NASH fibrosis (red) and in experimental NAFLD/NASH (C57 HFD vs AJ HFD; see Figs 4; black). The expression of the top predicted proteases (CAPN1/2 and MEP1A) was determined in liver samples from C57 and AJ mice (Panel B) by qPCR. The predicted substrates (**Panel C**) and GO Biological Process of these predicted substrates in the human samples (**Panel D**) are also shown. Summary data (**Panel B**) are box and whisker plots (n=4-8/group), which show the median (thick line), interquartile range (box) and range (whiskers). ^a^*P <*0.05 for effect of HFD, ^b^P <0.05 for effect of mouse strain by 2-way ANOVA using Tukey’s post hoc test.

We ran CausalMGM, a probabilistic graph modeling algorithm, to identify direct associations among peptidomic features and between peptidomic features and clinical data. The resulted graph encodes variables as nodes and direct connections between variables as (directed) edges. The first and second neighbors of the post-LT NASH fibrosis variable (≥F2) are presented in Figure 6A. The most informative variables for F2 are those in its Markov blanket (Figure 6A, circles): TYB4, VINC, FIBA, and NEUG. Besides Sex, no other clinical variable was found to be in the proximity of F2. Figure 6B shows the frequency distribution of the predicted proteases profiles for identified first- and second-degree neighbors in the Markov blanket identified by CausalMGM. In this subset of peptides, Capn1 and Mep1a were again the most predicted proteases similar to the degradome in toto (see Figures 1 and 5), although Mep1a was slightly enriched.

**Figure 6.**
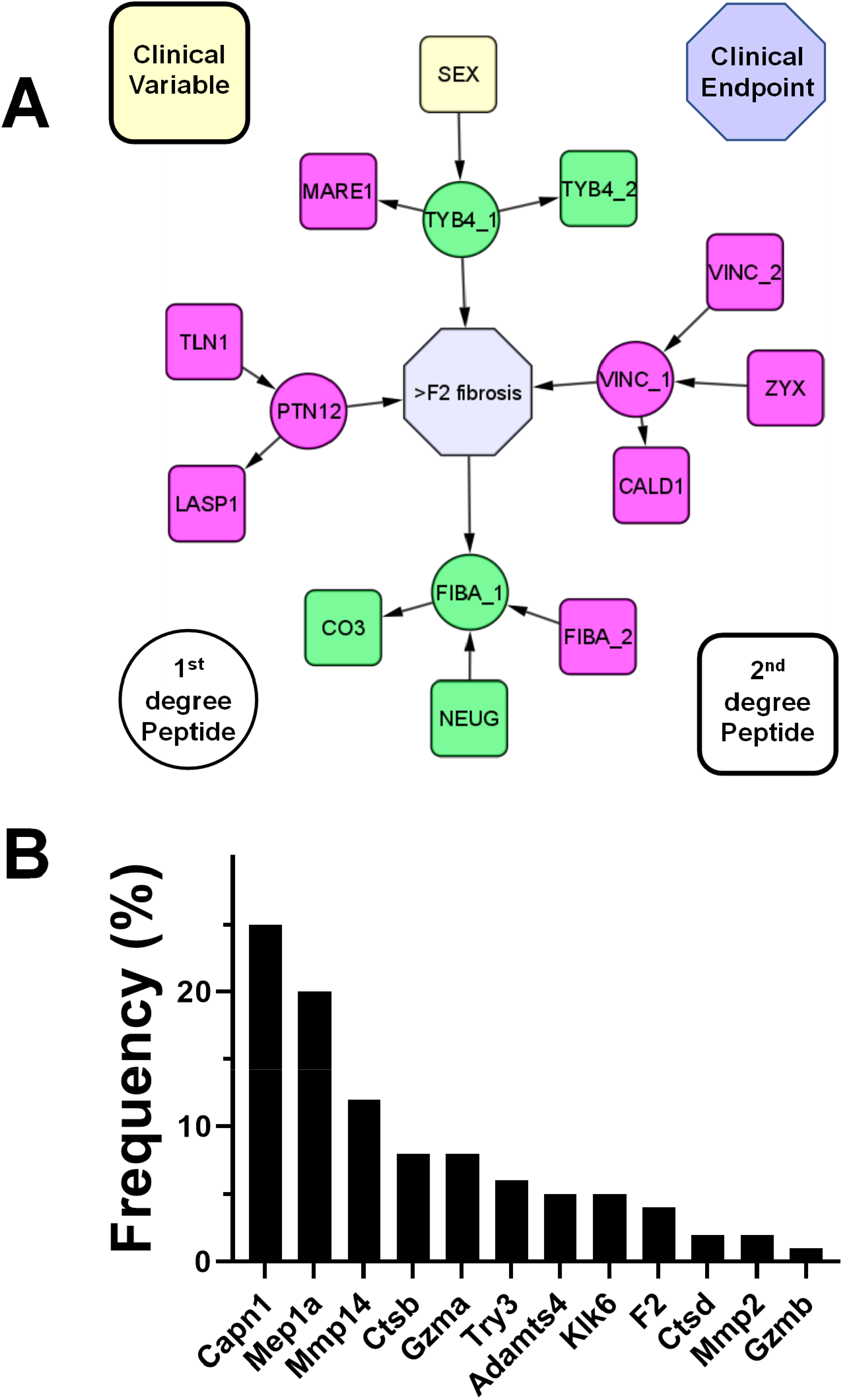
Probabilistic graphical modeling of predictive peptides. **Panel A** shows the probabilistic graph around the variable representing post-LT NASH fibrosis (≥F2). Only the first and second neighbors of F2 are depicted. The input dataset included all peptidome features and demographic and clinical variables. Based on this graph, the most informative variables for F2 are those in its Markov blanket (yellow nodes). **Panel B** shows frequency distribution of predicted proteases derived from the 1^st^ and 2^nd^ degree neighbors determined by probabilistic graphical modeling (Panel A). Colored nodes indicate fold-change of the peptides versus non-fibrosing controls (green increased, cyan decreased).

**Figure 7.**
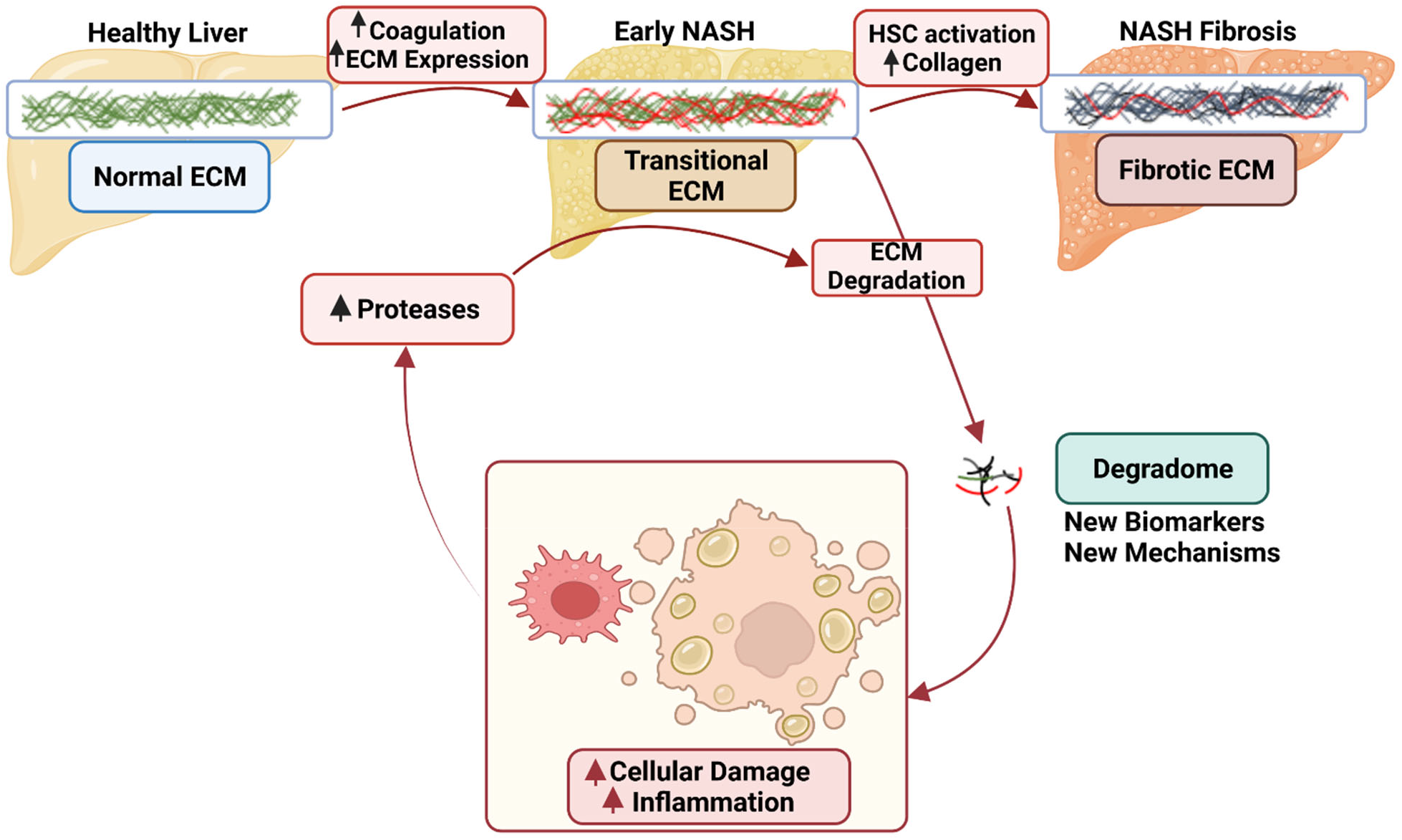
Working Hypothesis. Acute injury causes formation of a transitional ECM. This transitional ECM may contribute to injury and inflammation. The transitional ECM often resolves after removal of the insult and may contribute to the recovery from that insult. With continued injury, the transitional matrix may progress to a fibrotic matrix, which can also resolve under some conditions. Even during ECM accumulation in response to acute/chronic injury, there is a net increase in turnover of the matrisome, yielding degraded ECM products. We hypothesize that qualitative and quantitative changes to the “degradome” will serve as a biomarkers for outcome and mechanism in post-LT NASH fibrosis.

## Discussion

Liver transplantation often leads to an almost immediate improvement in the overall health and metabolism, as the patient transitions from a catabolic to anabolic state.^26^ Metabolic syndrome (MetS) is one of the most common post-transplant complications, with a prevalence of 44%-58% and, together with the immunosuppression, is considered the main risk factor for the development of cardiovascular disease in transplant recipients, which in turn accounts for 19%-42% of all deaths unrelated to the graft.^27^ As NAFLD is considered the hepatic manifestation of MetS, it is understandable that patients who undergo LT due to NASH-related cirrhosis frequently present recurrence of the disease after transplantation.^28^ However, it is hypothesized that immunosuppression also plays a crucial role in triggering or perpetuating the MetS.^29^ Thus, it is not surprising that not only recurrent, but also de novo forms of NAFLD, are frequent complications in patients who have undergone LT. Although there are approaches that detect those who have progressing liver fibrosis, including blood-based (e.g., FIB-4) and imaging-based (e.g., stiffness/elastography) noninvasive assessments, they are less reliable in post-LT patients and their accuracy is suboptimal for the F2 stage. What is needed is a relatively inexpensive test/score that better stratifies the risk for post-LT NASH fibrosis much earlier in disease progression, where interventive strategies to potentially halt disease progression would be more effective. Moreover, better understanding of drivers of post-LT NASH development could identify new mechanism-based interventive strategies.

During liver injury, perpetuation of the insult induces progressive accumulation of hepatic damage with incomplete remodeling, leading to increased ECM turnover as well as ECM accumulation.^30^ The end result of this dyshomeostasis is often fibrosis.^31^ Hepatic fibrogenesis is mediated by several converging pathways. These include (1) disruption of the normal ECM of the space of Disse because of hepatocyte damage and inflammatory infiltration; (2) release of reactive oxygen species and other fibrogenic/proinflammatory mediators (*e*.*g*., lysyl oxidases, transglutaminases, or MMPs); and (3) recruitment of immune cells, which in turn sustain fibrogenesis.^31^ Although hepatic fibrosis is considered almost synonymous with collagen accumulation,^11^ the qualitative and quantitative alterations to the hepatic ECM during fibrosis are much more diverse.^12^ Moreover, previous work has identified that hepatic remodeling occurs much sooner than fibrosis in disease progression.^30,32^ This understanding has let previous work to investigate the roles of non-collagen ECM components, as well as enzymes that metabolize them (e.g., MMPs) in the initiation and development of experimental and clinical fibrosis.^33,34^

Hypothesis-driven approaches are key to elucidating mechanism. However, the nature of the approach requires prior knowledge. Most research to date on hepatic fibrosis has focused on key collagenous ECM proteins identified histologically to be involved in fibrogenesis. In contrast, it is understood that remodeling of an organ is an orchestrated response not necessarily restricted to collagens or even ECM proteins. Discovery-based ‘omic techniques coupled with informatic analyses can yield new information and insight into disease progression and regression. In addition to synthesized peptides, the peptidome also contains fragments of proteins degraded by normal and/or abnormal processes (i.e., ‘degradome’) and shed into the surrounding biofluid space. Given the increased protein turnover associated with hepatic remodeling with disease, it was hypothesized that the degradome may serve as a new prognostic tool. These degradomic profiles are conditionally unique to the metabolic/pathologic status of the organism and reflect not only quantitative changes (i.e., more/less parent protein to degrade), but also qualitative changes (i.e., change in the pattern of protease digestion of parent proteins). The latter subset of the peptidome has generated key interest in some areas of human health as possible (surrogate) biomarkers for disease.

When the degradome was analyzed, an interesting pattern resolved. Specifically, there were distinct and significant differences in the pattern of the degradome in plasma from patients that later went on to develop at least significant (i.e., ≥F2) post-LT NASH fibrosis and those that did not (Figure 1). Unsupervised (heatmap and dendrogram) hierarchical clustering was able to separate the 2 groups well (Figure 1B). Moreover, supervised (OPLS-DA) discriminant analysis indicated that a fraction of the total peptide signal likely drives the assumed difference between the groups (Figure 1C). These data suggest that 3 months after LT and well before fibrosis was detectable with clinical assessment, that there are key differences in the degradome that can be observed.

Degradomic analysis allows for the integration of both quantitative (i.e., more/less parent protein degraded) and qualitative (i.e., change in the pattern of protease digestion of parent proteins) changes in the analysis (Figure 2). Increases the amount and number of ECM fragments previously associated with fibrosis development (e.g., collagens; Figure 2) was observed.^35^ However, one of the most robust family of degraded proteins that was enriched in “fibrosers” were associated with the coagulation cascade, with a particular enrichment of the fibrin(ogens) (e.g., FGA; Figure 2; supplemental table 3). Supervised discriminant analysis and machine learning approaches also ranked FGA peptides as important to predict post-LT NASH fibrosis development in human samples (Figures 1 and 6). Moreover, a similar enrichment of FGA fragments was observed when NASH-sensitive (C57Bl6/J) and -insensitive (AJ) mouse strains were compared (Figure 4). Although previous studies have investigated the potential role of fibrin(ogens) in the development of hepatic injury and fibrosis, the results are somewhat conflicting.^36,37^ In contrast, the turnover of fibrin(ogen) ECM has been heavily studied in other diseases of remodeling. For example, fibrin(ogen) ECM plays a key role in sub-cutaneous wound-healing.^38^ Since hepatic fibrosis and subcutaneous wound-healing qualitatively and quantitatively share many similarities, it is possible this this increase in fibrin(ogen) fragments represents an early detectable transition from provisional ECM to a collagenous ECM. Future studies should specifically investigate this possibility, especially given the use of anticoagulants in fibrotic liver disease and cirrhosis.

As mentioned in the Introduction, collagenous ECM accumulation has been the major focus in the field of hepatic fibrosis.^39^ Likewise, proteases involved in resolution of collagenous ECM (e.g., MMP2 and MMP9) have also received a sizeable amount of attention.^40^ The fragment sequence and putative cleavage site of the degradome may yield new information on proteases involved. The predicted protease that generated each peptide was therefore determined by in silico analysis of the cleavage site using the online open-source tool, Proteasix (www.proteasix.org; Figure 2).^25^ Although several proteases with established roles in fibrosis and/or resolution were identified (e.g., MMP2; Figure 2), several novel players were predicted by the algorithm. The most common predicted proteases in both humans and in our preclinical animal model were Capn1/2 and Mep1A (Figures 2 and 5). It should be noted that Capn1 and its isozyme Capn2 cannot be delineated by substrate or activity assays, and expression of both followed a similar pattern in liver during fibrosis in our preclinical model (Figure 5B). Moreover, there is significant overlap between Capn1/2 and Mep1a in predicted substrate targets (Figure 5C-5E). Capn1/2 and Mep1a are both cysteine proteases that cleave a myriad of protein substrates. Although they are generally localized intracellularly or on the plasma membrane, they can also be excreted in the active state.^41^ Capn1/2 and Mep1a activation has been identified as a key driver of inflammation in other diseases of inflammation and remodeling, such as atherosclerosis and cardiac disease.^41-44^ Interestingly, the predicted substrates for Capn1/2 and/or Mep1a activation not only included key ECM proteins [e.g., collagens and fibrin(ogens)], but also nuclear proteins, such as histones (Figure 5C). Few studies have investigated these proteases in hepatic injury.^45^ Moreover, recent work by this group showed that Capn2 is progressively induced in pretransplant NASH fibrosis severity.^46^ The mechanisms by which Capn2 may mediate these effects are unclear but work in other organ systems suggest a role in not only ECM metabolism, but also in organelle damage [e.g., ^41-44^]. These data suggest that the role of Capn1/2 and Mep1a in liver disease and fibrosis may be underappreciated at this time.

In conclusion, the identification of critical cleavage points of proteins is a novel and cutting-edge approach in the liver diseases field. The present study demonstrates the utility of degradomic approaches for development of novel biomarkers. The degradome characterization could lead to a better understanding of the specific pathological processes that trigger and perpetuate abnormal ECM remodeling that result in fibrosis and cirrhosis. Degradomic peptide patterns may also play a role in predicting early liver fibrosis onset in the setting of post-LT NAFLD/NASH and, consequently, LT outcomes. The identified protein products could become potentially targetable molecules to develop new therapies against the progression of liver fibrosis. Further prospective and etiology-controlled studies are needed to confirm the results.

## Methods

### Biobanked patient samples

This was a retrospective analysis that compared 12 post-LT patients (all living donor recipients) that developed significant biopsy-proven NASH fibrosis to 10 age/sex/weight-matched patients that did not (see Table I). For the purpose of this study, we defined significant NASH fibrosis as the development of ≥F2 fibrosis (Brunt-Kleiner score) within 3 years post-LT. According to an institutional research protocol, LT candidates were approached by the time of evaluation (i.e., pre-LT) by research coordinators and asked to participate in UPMC LT Biobanking Protocol. Following consent, a blood sample was drawn 3 months after LT along with a clinic visit’s laboratories and plasma was aliquoted and stored at – 80°C. Those candidates successfully achieving transplantation continued to participate in the post-LT period contributing with specimens for biobanking. Samples and relevant deidentified clinical data (e.g., liver chemistry and biopsy results) were collected from recipient pre-LT, donor pre-LT (for living donor LT), 3, 6 and 12 months post-LT, and annually thereafter. The 3-month sample served as a “baseline” for each patient and is reported to remove influences of donor and transplant.^3^

**Table I.** Baseline characteristics of LT patients.

|  | ***Non-progressors*** | ***Progressors*** | ***P-value*** |
| --- | --- | --- | --- |
| *Age* | 62±7 | 54±14 | 0.12 |
| *M/F* | 5/5 | 6/6 | 1.0 |
| *Etiology* | NASH 4<br>ALD 2<br>HCV 1<br>Autoimmune 0<br>Other 1<br>Cryptogenic 2 | NASH 1<br>ALD 1<br>HCV 2<br>Autoimmune 4<br>Other 1<br>Cryptogenic 3 | 0.16 |
| *BMI* | 30 ± 5 | 28 ± 5 | 0.39 |
| *Bariatric surgery* | 0% | 8% | 0.54 |

All transplant recipients were from living donors (LDLT). There were no differences in key baseline characteristics between transplant recipients that developed fibrosis (F≥2 progressors) and those who did not. There were no patients with diabetes mellitus at baseline. Among the group of progressors, 4 recipients (33%) ultimately developed cirrhosis.

### Clinical endpoints

De-identified post-LT records were obtained through an honest broker. Per protocol, our program performs allograft biopsies and liver imaging at 1 year after living donor LT, which are mandated in addition to clinically-indicated investigations. Imaging-based non-invasive liver disease assessment is also obtained on a yearly basis, including magnetic resonance elastography and proton-derived fat fraction. Post-LT NAFLD was defined as steatosis ≥5% on liver biopsy or as a proton-density fat fraction ≥6.4%, whereas NASH was defined as the presence of steatosis plus steatohepatitis or a perivenular pattern of fibrosis suggestive of NASH per pathology reports. The primary endpoint from our clinical cohort was progression to at least significant fibrosis (F≥2), defined on the basis of liver biopsy, magnetic resonance elastography ≥3.1 kPa or obvious imaging features of cirrhosis plus incidental features and clinical complications from portal hypertension (splenomegaly excluded). For recurrent cirrhosis, a threshold of 4.7 kPa was utilized for magnetic resonance elastography.

### Animals and Treatments

Previous studies have indicated that there are strain differences in mice in the sensitivity of the liver to damage caused by an obesogenic diet.^47^ Specifically, although AJ and C57Bl/6J mice both develop insulin resistance when fed HFD, the latter strain develops much more robust liver disease.^23,24,48,49^ Accordingly, 8 weeks old, male AJ and C57Bl/6J mice purchased from Jackson Laboratory (Bar Harbor, ME) were housed in a pathogen-free barrier facility accredited by the Association for Assessment and Accreditation of Laboratory Animal Care, and procedures were approved by the local Institutional Animal Care and Use Committee. Animals were allowed standard laboratory chow and water *ad libitum*. Mice were fed either a low-fat control diet (LFD: 13% saturated fat) or a ‘Western’-style high-fat, high-fructose diet (HFD: 42% saturated fat) for 12 weeks as described previously.^50,51^ Animals were weighed on a weekly basis to check growth and food consumption was monitored. Mice were fasted for 4 hours prior to sacrifice. At the time of sacrifice, animals were anesthetized with ketamine/xylazine (100/15 mg/kg, i.p.). Blood was collected from the vena cava just prior to sacrifice by exsanguination and citrated plasma was stored at -80°C for further analysis. Portions of liver tissue were frozen immediately in liquid nitrogen, while others were fixed in 10% neutral buffered formalin or embedded in frozen specimen medium (Tissue-Tek OCT compound, Sakura Finetek, Torrance, CA) for subsequent sectioning and mounting on microscope slides.

### RNA Isolation and Real-Time RT-PCR

RNA extraction and real-time RT-PCR were performed as described previously ^51^. RNA was extracted immediately following sacrifice from fresh liver samples using RNA Stat60 (Tel-Test, Ambion, Austin, TX) and chloroform. RNA concentrations were determined spectrophotometrically and 1µg of total RNA was reverse transcribed using the QuantiTect Reverse Transcription Kit (Qiagen,Valencia, CA). Real-time RT-PCR was performed using a StepOne real time PCR system (Thermo Fisher Scientific, Grand Island, NY) using the Taqman Universal PCR Master Mix (Life Technologies, Carlsbad, CA). Primers and probes were ordered as commercially available kits (Thermo Fisher Scientific, Grand Island, NY; see Supplemental Table 1). The comparative C_T_ method was used to determine fold differences between the target genes and an endogenous reference gene (18S). Results were reported as copy number (2^-ΔCT^) or fold naïve control (FC; 2^-ΔΔ*C*T^).

### Biochemical assays and histology

Plasma levels of Aspartate transaminase (AST) and alanine amino transferase (ALT) were determined spectrophotometrically using standard kits (Thermo Fisher Scientific, Waltham, MA), as described previously.^32^ Formalin-fixed, paraffin embedded sections were cut at 5 μm and mounted on glass slides. Deparaffinized sections stained with hematoxylin and eosin and pathology was assessed in a blinded manner. ECM accumulation in liver sections was determined by staining with Sirius red/fast green and were visualized via brightfield.^52^

### Plasma peptide purification and high resolution LC-MS/MS analysis

1 volume of PBS and 0.025 volumes of 10X iRT standards (Biognosis Inc, Beverly, MA) were added to 1 volume of plasma. 20% w/v TCA was mixed into each sample on ice to a final concentration of 10% w/v. Samples were then incubated for 1 hour at 4°C. The precipitate was pelleted by centrifugation at 16,000 × G for 10 min at 4°C; pellets were discarded. The centrifugation was then repeated with the supernatant samples, followed by discard of the second pellets. Samples were then desalted and concentrated using solid phase extraction (Waters Oasis HLB µElution 30 µm plate, part no. 186001828BA) as described.^52,53^ Solid phase extraction eluates were dried in a SpeedVac, and dried peptides were re-suspended in 2% v/v acetonitrile/ 0.1% v/v formic acid to yield an iRT concentration of 1X (assuming complete recovery). 1µL of each sample was separated using an Acclaim PepMap 100 75µm x 2cm, nanoViper (C18, 3µm, 100Å) trap, and an Acclaim PepMap RSLC 50µm x 15cm, nanoViper (C18, 2µm, 100Å) separating column (ThermoFisher Scientific, Waltham, MA, USA) for mouse degradome samples or an Acclaim PepMap RSLC 75µm x 50cm, nanoViper (C18, 2µm, 100Å) separating column (ThermoFisher) for human degradome samples. Peptides were eluted under a 60min 2%-37% acetonitrile gradient for mouse degradome (2%-45% for human degradome) and transferred by Nanospray Flex nanoelectrospray source (ThermoFisher) into an Orbitrap Elite - ETD mass spectrometer (ThermoFisher) using an Nth Order Double Play (Xcalibur v2.2, ThermoFisher) with FTMS MS1 scans (240,000 resolution) collected from 300-2000m/z and ITMS MS2 scans collected on up to twenty peaks having a minimum signal threshold of 5,000 counts from the MS1 scan event.

### Peptidomic Data Analysis

RAW files were searched in Peaks Studio X (Bioinformatics Solutions Inc., Waterloo, ON, Canada) using the Denovo, PeaksDB, PeaksPTM, and Label Free Q algorithms. The UniprotKB reviewed reference proteome canonical and isoform Homo sapiens sequences (Proteome ID UP000005640) version 11/30/2020 was used for the human degradome samples and the Mus musculus sequences (Proteome ID UP000000589) version 7/22/2021 for the mouse degradome samples. Search parameters included: variable methionine and proline oxidation (+15.9949 Da), no enzyme specified, 15 ppm precursor error for MS1 Orbitrap FTMS data, 0.5 Da error for MS2 data sets, and selected common PTMs in the PeaksPTM algorithm. The peptide, feature, and protein-peptide csv files were exported from the Label Free Q result for statistical tests in Microsoft Excel 2016 and R v3.5.0. High confidence peptide assignments (Peaks Studio X criteria –logP scores with FDR threshold 1%) were exported into comma separated values files (.csv) for upload and analysis by the Proteasix (http://www.proteasix.org) algorithm using the ‘Peptide-centric’ prediction tool based on curated, known and observed cleavage events enabling assignment of protease identities from the MEROPS database as described previously.^54^ The positive predictive value (PPV) cut-off for the prediction algorithm output was set to 80%. Protein-protein interaction network analysis of regulated proteomic data sets (q-value <0.05) was performed using Search Tool for the Retrieval of Interacting Genes/Proteins, STRING v11,^19^ with the highest confidence score (0.900). The resultant matrix of both Proteasix and STRING analysis were visualized using Cytoscape v3.9.1. Node sizes of the predicted proteases represented the relative frequency with which the top 15 proteases were predicted to mediate the observed cleavage (0.2-27%). Node sizes of the peptides represented the relative number of unique peptides (1-56) identified from each parent protein. Node color of the peptides represented the median Log2FC vs non-fibrosers for all peptides derived from that parent protein.

### Probabilistic Graphical Modeling

We normalized the peptidomic data using nonparanormal (npn) transformation,^56^ and combined the normalized data with clinical variables including sex, age, fibrosis stage (F2-F4), NAFLD/NASH diagnosis, rejection and years follow-up to create a data matrix with mixed data types. We ran CausalMGM,^57^ with the Fast Greedy Equivalence Search (FGES) algorithm (penalty discount = 6) to construct a probabilistic graph and identify variables in the Markov blanket of F2 fibrosis.

## Data Sharing

Proteomic files will be deposited in MassIVE (http://massive.ucsd.edu/) under a study entitled “Post-liver transplant plasma degradome reflects later development of severe NASH fibrosis”. Data will include (A) the primary data files (.RAW), (B) peak list files (.mzML), (C) sample key, (D) the sequence databases (mouse and human UniprotKB reviewed reference proteoms), and (E) excel files containing Peaksdb results for de novo peptide sequence assignment. Shared data will be released from private embargo for public access upon acceptance for publication.

## Abbreviations

ALT: alanine aminotransferase
AST: aspartate aminotransferase
BW: body weight
CVD: cardiovascular disease
ECM: extracellular matrix
GO: gene ontology
HFD: high-fat diet
HOMA-IR: homeostatic model assessment for insulin resistance
KEGG: Kyoto encyclopedia of genes and genomes
LC-MS/MS: liquid chromatography, tandem mass-spectroscopy
LFD: low-fat diet
Log2FC: fold-change (log2)
LT: liver transplantation
MetS: metabolic syndrome
MMP: matrix metalloproteinases
MS: mass spectrometry
NAFLD: non-alcohol-related fatty liver disease
NASH: non-alcoholic steatohepatitis
OPLS-DA: orthogonal partial least squares discriminant analysis.

## Acknowledgements

Supported, in part, by grants from NIH (R01 DK130294, R01 AA021978, P20 GM113226, P30 DK120531).

## Author Contributions

JL: Visualization, Investigation, Validation, Formal Analysis, Writing-Original Draft. TS: Investigation, Validation, Formal Analysis. MH-T: Writing-Original Draft. JIB: Investigation, Validation, Formal Analysis. KS: Visualization, Formal Analysis, Writing-Original Draft. PVB: Visualization, Conceptualization, Supervision, Writing-Original Draft, Funding acquisition, Resources. DW: Investigation, Validation. AH: Formal Analysis, Resources. MLM: Visualization, Conceptualization, Supervision, Writing-Original Draft, Funding acquisition, Resources. AD-R: Visualization, Conceptualization, Supervision, Writing-Original Draft, Funding acquisition, Resources. GEA: Project Administration, Visualization, Conceptualization, Investigation, Supervision, Writing-Review and Editing, Funding acquisition, Resources

## Data availability statement

Proteomic files will be deposited in MassIVE (http://massive.ucsd.edu/) under a study entitled “Post-liver transplant plasma degradome reflects later development of severe NASH fibrosis”. Data will include (A) the primary data files (.RAW), (B) peak list files (.mzML), (C) sample key, (D) the sequence databases (mouse and human UniprotKB reviewed reference proteoms), and (E) excel files containing Peaksdb results for de novo peptide sequence assignment. Shared data will be released from private embargo for public access upon acceptance for publication. All other d ata will be made available on request.

## Additional Information

The authors declare that they have no known competing financial interests or personal relationships that could have appeared to influence the work reported in this paper.

