## supplemental materials for "The plasma degradome reflects later development of NASH fibrosis after liver transplant"

Running title: The degradome in post-LT NASH fibrosis.

Key Words: Extracellular matrix, remodeling, liver transplantation outcomes, translational study

Send all correspondence to: Gavin E. Arteel, PhD, FAASLD
Thomas E. Starzl Biomedical Science Tower
West 1143
200 Lothrop Street
Pittsburgh, PA 15213


**Abbreviations**: ALT, alanine aminotransferase; AST, aspartate aminotransferase; BW; body weight; CVD, cardiovascular disease; ECM, extracellular matrix; GO, gene ontology; HFD, high-fat diet; HOMA-IR, homeostatic model assessment for insulin resistance; KEGG, Kyoto encyclopedia of genes and genomes; LC-MS/MS, liquid chromatography, tandem mass-spectroscopy; LFD, low-fat diet; Log2FC, fold-change (log2); LT, liver transplantation; MetS, metabolic syndrome; MMP, matrix metalloproteinases; MS, mass spectrometry; NAFLD, non-alcohol-related fatty liver disease; NASH, non-alcoholic steatohepatitis; OPLS-DA, orthogonal partial least squares discriminant analysis.

**Supplemental Methods**

Information on rtPCR primers and probes, drugs and chemical assays and antibodies used in this study are summarized in supplemental Tables 1-2.

Biochemical assays and histology. Plasma levels of Aspartate transaminase (AST) and alanine amino transferase (ALT) were determined spectrophotometrically using standard kits (Thermo Fisher Scientific, Waltham, MA), as described previously.[1] Formalin-fixed, paraffin embedded sections were cut at 5 µm and mounted on glass slides. Deparaffinized sections stained with hematoxylin and eosin and pathology was assessed in a blinded manner. ECM accumulation in liver sections was determined by staining with Sirius red/fast green and were visualized via brightfield analysis.[2] Hepatic lipids (NEFA, TG and TC) were determined spectrophotometrically, as described previously.[3] Fasting plasma glucose and insulin levels were determined using commercially-available kits (Jiang need information). HOMA-IR was calculated using the following equation: fasting insulin (ng/mL)×fasting blood glucose (mg/dL)/405.[4]

RNA Isolation and Real-Time RT-PCR RNA extraction and real-time RT-PCR were performed as described previously.[5] RNA was extracted immediately following sacrifice from fresh liver samples using RNA Stat60 (Tel-Test, Ambion, Austin, TX) and chloroform. RNA concentrations were determined spectrophotometrically and 1µg of total RNA was reverse transcribed using the QuantiTect Reverse Transcription Kit (Qiagen,Valencia, CA). Real-time RT-PCR was performed using a StepOne real time PCR system (Thermo Fisher Scientific, Grand Island, NY) using the Taqman Universal PCR Master Mix (Life Technologies, Carlsbad, CA). Primers and probes were ordered as commercially available kits (Thermo Fisher

Scientific, Grand Island, NY). Primer sequences from Taqman Gene Expression Assays (Applied Biosystems, Foster City, CA) were as follows: Collagen type I alpha 1 (Col1α1), Actin Alpha 2, Smooth Muscle (ACTA2), Transforming Growth Factor Beta 1 (TGFB1), Prolyl 4-Hydroxylase, Transmembrane (P4htm), protein-lysine 6-oxidase (LOX1,2), MMP- 2, 8, 9, 12, 13, and 14. The comparative $C_T$ method was used to determine fold differences between the target genes and an endogenous reference gene (18S). Results were reported as copy number $2^{-\Delta CT}$.

### Supplemental Results

Supplemental Table 4 summarizes results of cluster analysis of significantly increased peptides by StringDB, which is also shown graphically in Figure 5. Supplemental Table 5 lists all peptides significantly increased in mouse plasma 1D and/or 28D after cessation of $CCl_4$, as described in Experimental Procedures, which is shown graphically in Figure 5. Supplemental Table 6 summarizes feeding, growth and lipid accumulation caused by HFD in C57Bl6/J and AJ mice (see also Figure 3). Supplemental Figure 1 compares mRNA expression of key genes involved in hepatic inflammation and fibrosis in C57Bl6/J and AJ mice fed HFD or LFD for 12 weeks (see also Figure 3).

**Supplemental Table 1-Product information for primers used in RT-PCR**

| Gene name | Supplier | Cat. No. |
|---|---|---|
| *18s* | Thermofisher | Mm03928990_g1 |
| *Col1a1* | Thermofisher | Mm00801666_g1 |
| *Capn1* | Thermofisher | Mm00482964_m1 |
| *Capn2* | Thermofisher | Mm00486669_m1 |
| *Mep1a* | Thermofisher | Mm00484970_m1 |
| *Tnfa* | Thermofisher | Mm00443258_m1 |
| *Il10* | Thermofisher | Mm01288386_m1 |
| *Serpine1* | Thermofisher | Mm00435858_m1 |
| *Tgfb1* | Thermofisher | Mm01178820_m1 |
| *Acta2* | Thermofisher | Mm01546133_m1 |
| *Lox1* | Thermofisher | Mm00454582_m1 |
| *Lox2* | Thermofisher | Mm01325281_m1 |
| *Mmp2* | Thermofisher | Mm00439498_m1 |
| *Mmp9* | Thermofisher | Mm00442991_m1 |

**Supplemental Table 2- Supplies and chemical assays used in this study**

| Category | Name | Supplier | Cat No. |
|---|---|---|---|
| *Diets* | Western HFD | Envigo | TD.88137 |
| | Control LFD | Envigo | TD.06416 |
| *Chemical assays* | AST assay kit | Thermofisher | TR70121 |
| | ALT assay kit | Thermofisher | TR71121 |
| | Free fatty acid kit | Sigma | MAK044-1KT |
| | Cholesterol assay kit | Thermofisher | TR13421 |
| | Triglyceride assay kit | Thermofisher | TR22421 |
| | Insulin ELISA kit | ALPCO | 80-INSMR-CH10 |
| | Glucose strip | Bayer HealthCare | 7090G |

**Supplemental Table 3-Plasma peptides changed in post-LT fibrosis**

| Protein | Accession | Start | End | Sequence | Log2FC | -Log10PV |
|---|---|---|---|---|---|---|
| A1AG2 | P19652 | 134 | 145 | LDDEKNWGLSFY | -1.84 | 2.73 |
| A2AP | P08697 | 42 | 53 | EQVSPLTLLKLG | -3.34 | 8.75 |
| ACTA | P62736 | 194 | 202 | ILTERGYSF | -1.54 | 1.33 |
| ACTB | P60709 | 2 | 15 | D(+42.01)DDIAALVVD(+21.98)NGSG | 4.66 | 2.40 |
| | | 2 | 11 | D(+42.01)DDIAALVVD | 1.39 | 2.58 |
| | | 20 | 36 | GFAGDDAPRAVFPSIVG | 1.15 | 2.76 |
| | | 24 | 36 | DDAPRAVFPSIVG | -3.96 | 5.12 |
| | | 49 | 67 | Q(-17.03)KDSYVGDEAQSKRGILTL | -1.87 | 3.00 |
| | | 54 | 66 | VGDEAQSKRGILT | 2.59 | 2.48 |
| | | 99 | 113 | EEHPVLLTEAPLNPK | -2.04 | 1.88 |
| | | 162 | 174 | TVPIYEGYALPHA | -1.66 | 1.53 |
| ANDR | P10275 | 447 | 480 | YGPCGGGGGGGGGGGGGGGGGGGGGGGEAGAVAP | 1.63 | 1.37 |
| | | 455 | 477 | GGGGGGGGGGGGGGGGGGGEAGA | -4.66 | 9.73 |
| AP2B1 | P63010 | 678 | 694 | APSPTPAVVSSGLNDLF | -1.96 | 3.55 |
| APOC3 | P02656 | 82 | 99 | SEFWDLDPEVRPTSAVAA | 4.94 | 5.45 |
| APOF | Q13790 | 314 | 326 | SYDLDPGAGSLEI | -3.36 | 3.70 |
| BIN2 | Q9UBW5 | 263 | 272 | SPPVRTATVS | 2.38 | 4.42 |
| | | 536 | 565 | QLQVSMVPENNNLTAPEPQEEVSTSENPQL | 3.71 | 7.66 |
| | | 541 | 565 | MVPENNNLTAPEPQEEVSTSENPQL | -1.11 | 2.40 |
| C1QB | P02746 | 83 | 101 | PGNPGKVGPKGPMGPKGGP | 6.19 | 11.83 |
| CALD1 | Q05682 | 119 | 130 | KEFDPTITDASL | -1.48 | 2.24 |
| | | 120 | 134 | EFDPTITDASLSLPS | -3.78 | 9.02 |
| | | 716 | 722 | WEKGNVF | 2.61 | 5.77 |
| CASS4 | Q9NQ75 | 164 | 180 | ASLPTLPSQVYDVPTQH | 3.88 | 7.11 |
| CAVN2 | O95810 | 21 | 31 | EKPSSPSPM(+15.99)PS | 3.36 | 2.45 |
| | | 234 | 243 | SSLKKVDSLK | 1.32 | 1.42 |
| | | 235 | 247 | SLKKVDSLKKAFS | 6.61 | 11.83 |
| | | 288 | 294 | SPFKVSP | 2.42 | 1.82 |
| CD99 | P14209 | 175 | 184 | EPAVQRTLLE | 1.11 | 1.37 |
| CFAB | P00751 | 239 | 257 | TETIEGVDAEDGHGPGEQQ | 2.86 | 7.30 |
| CMGA | P10645 | 99 | 131 | GFEDELSEVLENQSSQAELKEAVEEPSSKDVME | -4.73 | 10.05 |
| CMIP | Q8IY22 | 720 | 732 | ETPVTDAGLLALS | 2.07 | 1.98 |
| CNN2 | Q99439 | 2 | 15 | S(+42.01)S(+79.97)TQFNKGPSYGLS | 1.92 | 2.23 |
| CNST | Q6PJW8 | 320 | 334 | TESSKESQHTVEPLG | 3.14 | 2.67 |
| CO1A1 | P02452 | 371 | 397 | GEPGPPGPAGAAGPAGNPGADGQPGAK | 1.54 | 2.18 |
| | | 546 | 558 | SP(+15.99)GPDGKTGPP(+15.99)GP | 2.29 | 6.16 |
| | | 818 | 844 | GADGQPGAKGEP(+15.99)GDAGAKGDAGPP(+15.99)GPA | 5.70 | 10.98 |
| | | 819 | 844 | ADGQPGAKGEP(+15.99)GDAGAKGDAGPP(+15.99)GPA | 2.39 | 3.92 |
| | | 820 | 843 | DGQP(+15.99)GAKGEPGDAGAKGDAGPPGP(+15.99) | 7.13 | 11.17 |
| | | 1151 | 1203 | GKDGLNGLPGPIGPPGPRGRTGDAGPVGPPGPPGPPGPPGPPSAGFDFSFLPQ | -10.05 | 20.03 |
| CO3 | P01024 | 955 | 982 | EGVQKEDIPPADLSDQVPDTESETRILL | 2.31 | 2.04 |

| Protein | Accession | Start | End | Sequence | Log2FC | -Log10PV |
|---|---|---|---|---|---|---|
| | | 969 | 987 | DQVPDTESETRILLQGTPV | 3.02 | 6.79 |
| | | 1321 | 1343 | SEETKENEGFTVTAEGKGQGTLS | 7.10 | 6.49 |
| | | 1321 | 1341 | SEETKENEGFTVTAEGKGQGT | 5.83 | 11.83 |
| | | 1322 | 1336 | EETKENEGFTVTAEG | 4.56 | 3.06 |
| CO3A1 | P02461 | 500 | 510 | P(+15.99)GEKGPAGERG | 2.54 | 2.75 |
| | | 1151 | 1166 | DGTS(-18.01)GHPGPIGPP(+15.99)GPR(+28.03) | 8.60 | 8.63 |
| CO4A | P0C0L4 | 957 | 979 | TLEIPGNSDPNM(+15.99)IPDGDFNSYVR | 2.73 | 6.41 |
| | | 1337 | 1349 | NGFKSHALQLNNR | 1.77 | 1.54 |
| | | 1429 | 1449 | DDPDAPLQPVTPLQLFEGRR(+0.98)N | 3.43 | 5.28 |
| | | 1429 | 1439 | DDPDAPLQPVT(-18.01) | 5.04 | 8.26 |
| | | 1431 | 1443 | PDAPLQPVTPLQL | 1.05 | 2.30 |
| CO4A2 | P08572 | 365 | 386 | PGFPGAQGEPGSQGEPGDPGLP | 1.96 | 4.04 |
| CO7A1 | Q02388 | 1783 | 1797 | LDGKPGAAGPSGPNG | -4.44 | 5.53 |
| COMA1 | Q8NFW1 | 911 | 922 | APGAAGNPGAPG | 1.31 | 1.57 |
| CPNS1 | P04632 | 39 | 60 | GGGGGGGGGGGGGGGGGT(-18.01)AMR(+21.98) | -2.64 | 8.09 |
| CRKL | P46109 | 214 | 239 | TPLPAVSGSPGAAITPLPSTQNGPVF | 1.03 | 2.30 |
| DOK3 | Q7L591 | 358 | 369 | AEPGPQSLPLLL | 2.79 | 2.81 |
| DREB | Q16643 | 33 | 61 | TYEDGSDDLKLAASGEGGLQELSGHFENQ | 2.04 | 3.59 |
| | | 467 | 490 | NVPPAATSLIDLWPGNGEGASTLQ | 1.92 | 1.54 |
| | | 468 | 490 | VPPAATSLIDLWPGN(+.98)GEGASTLQ | -1.16 | 1.65 |
| | | 468 | 490 | VPPAATSLIDLWPGNGEGASTLQ | 1.11 | 1.62 |
| | | 473 | 490 | TSLIDLWPGNGEGASTLQ | -2.70 | 4.60 |
| EPN4 | Q14677 | 322 | 338 | SSGDLVDLFDGTSQSTG | -1.58 | 4.10 |
| ESYT2 | A0FGR8 | 311 | 319 | MPLVGALSI | -3.28 | 3.14 |
| FBLN3 | Q12805 | 112 | 131 | VLPGGGFVASAAAVAGPEMQ | -1.52 | 1.51 |
| FHOD1 | Q9Y613 | 365 | 389 | RRSLEGGGCPARAPEPGPTGPASPV | 6.47 | 8.09 |
| FIBA | P02671 | 20 | 35 | ADS(+79.97)GEGD(-18.01)FLAEGGGVR | -3.47 | 9.97 |
| | | 20 | 34 | AD(-18.01)SGEGDFLAEGGGV | 5.10 | 6.80 |
| | | 24 | 34 | EGDFLAEGGGV | -2.96 | 6.53 |
| | | 26 | 35 | DFLAEGGGVR | -1.43 | 2.19 |
| | | 29 | 38 | AEGGGVRGPR | 3.65 | 6.43 |
| | | 229 | 238 | VPDLVPGNFK | 2.22 | 3.84 |
| | | 231 | 237 | DLVPGNF | 3.00 | 2.63 |
| | | 260 | 269 | ELERPGGNEI | -2.08 | 1.68 |
| | | 272 | 296 | GGSTSYGTGSETESPRNPSSAGSWN | 5.61 | 7.76 |
| | | 300 | 332 | SGPGSTGNRNPGSSGTGGTATWKPGSSGPGSTG | -1.53 | 5.44 |
| | | 303 | 328 | GSTGN(+0.98)RNPGSSGTGGTATWKPGSSGP | -4.50 | 5.36 |
| | | 330 | 361 | S(+27.99)TGSWNSGSSGTGSTGNQNPGSPRPGSTGTWN | -2.70 | 3.25 |
| | | 380 | 407 | SGSTGQWHSESGSFRPDSPGSGNARPNN | 6.45 | 10.18 |
| | | 380 | 406 | SGSTGQWHSESGSFRPDSPGSGNARPN.N | 6.29 | 10.61 |
| | | 392 | 424 | SFRPDSPGSGNARPNNPDWGTFEEVSGNVSPGT | -1.18 | 2.59 |
| | | 406 | 425 | NNPDWGTFEEVSGNVSPGTR | -1.46 | 1.86 |
| | | 426 | 443 | REYHTEKLVTSKGDKELR | 1.15 | 1.68 |

| Protein | Accession | Start | End | Sequence | Log2FC | -Log10PV |
|---|---|---|---|---|---|---|
| | | 430 | 441 | TEKLVTSKGD(-18.01)KE | -1.52 | 1.73 |
| | | 521 | 536 | DTASTGKTFPGFFSPM(+15.99) | 1.34 | 1.67 |
| | | 525 | 535 | TGKTFPGFFSP | 5.30 | 8.33 |
| | | 537 | 557 | LGEFVSETESRGSESGIFTNT | 1.48 | 2.83 |
| | | 538 | 557 | GEFVSETESRGSESGIFTNT | -1.41 | 3.36 |
| | | 542 | 600 | SETESRGSESGIFTNTKESSSHHPGIAEFPSRGKSSSYSKQFTSSTSYNRGDSTFESKS | 7.07 | 7.25 |
| | | 542 | 598 | SETESRGSESGIFTNTKESSSHHPGIAEFPSRGKSSSYSKQFTSSTSYNRGDSTFES | 6.79 | 7.76 |
| | | 542 | 587 | SETESRGSESGIFTNTKESSSHHPGIAEFPSRGKSSSYSKQFTSST | 7.87 | 4.15 |
| | | 542 | 582 | SETESRGSESGIFTNTKESSSHHPGIAEFPSRGKSSSYSKQ | 7.09 | 6.42 |
| | | 542 | 577 | SETESRGSESGIFTNTKESSSHHPGIAEFPSRGKSS | 7.61 | 4.89 |
| | | 542 | 576 | SETESRGSESGIFTNTKESSSHHPGIAEFPSRGKS | 2.12 | 2.82 |
| | | 542 | 574 | SETESRGSESGIFTNTKESSSHHPGIAEFPSRG | 5.63 | 2.87 |
| | | 542 | 570 | SETESRGSESGIFTNTKESSS(-18.01)HHPGIAEF | 6.69 | 8.36 |
| | | 542 | 566 | SETESRGSESGIFTNTKESSSHHPG | 3.87 | 2.77 |
| | | 542 | 564 | SETESRGSESGIFTNTKESSSHH | 5.81 | 11.83 |
| | | 542 | 562 | SETESRGSESGIFTNTKESSS | 2.82 | 8.33 |
| | | 542 | 556 | SETESRGSESGIFTN | 6.05 | 11.56 |
| | | 543 | 556 | ETESRGSESGIFTN | -3.15 | 2.70 |
| | | 543 | 555 | ETESRGSESGIFT | 4.64 | 4.89 |
| | | 551 | 566 | SGIFTNTKESSSHHPG(+15.99) | 3.24 | 2.03 |
| | | 552 | 568 | GIFTNTKESSSHHP(+15.99)GIA | 2.40 | 4.80 |
| | | 559 | 578 | ESSSHHPGIAEFPSRGKSSS | 2.27 | 2.21 |
| | | 576 | 596 | SSSYSKQFTSSTSYNRGD(-18.01)STF | 1.71 | 1.97 |
| | | 582 | 604 | Q(-17.03)FTSSTSYNRGDSTFESKSYKMA | 4.85 | 9.13 |
| | | 582 | 598 | QFTSSTSYNRGDSTFES | 3.16 | 6.27 |
| | | 584 | 596 | TSSTSYNRGDSTF | 3.00 | 2.49 |
| | | 586 | 603 | STSYNRGDSTFESKSYKM(+15.99) | -2.81 | 5.61 |
| | | 590 | 601 | NR(+0.98)GDSTFESKSY | 5.84 | 11.83 |
| | | 597 | 619 | ESKSYKM(+15.99)ADEAGSEADHEGTHST | 5.27 | 7.94 |
| | | 600 | 629 | SYKMADEAGSEADHEGTHSTKRGHAKSRPV | 4.96 | 3.75 |
| | | 602 | 624 | KMADEAGSEADHEGTHSTKRGHA | 5.62 | 10.98 |
| | | 602 | 622 | KM(+15.99)ADEAGSEADHEGTHSTKRG | 2.26 | 2.40 |
| | | 603 | 629 | MADEAGSEADHEGTHSTKRGHAKSRPV | -3.32 | 3.51 |
| | | 605 | 629 | DEAGSEADHEGTHSTKR(+0.98)GHAKSRPV | 3.33 | 2.48 |
| | | 605 | 628 | DEAGSEADHEGTHSTKR(+0.98)GHAKSRP | 1.50 | 2.28 |
| | | 607 | 629 | AGSEADHEGTHSTKR(+0.98)GHAKSRPV | 2.60 | 2.05 |
| | | 607 | 619 | AGSEADHEGTHST | 3.85 | 5.83 |
| | | 609 | 624 | SEADHEGTHSTKRGHA | 2.68 | 4.49 |
| | | 610 | 624 | EADHEGTHSTKRGHA | 2.32 | 1.90 |
| FIBB | P02675 | 31 | 50 | Q(-17.03)GVNDNEEGFFSAR(+0.98)GHRPLD | 5.61 | 10.31 |
| | | 31 | 50 | QGVNDNEEGFFS(-18.01)AR(+0.98)GHRPLD | 7.86 | 10.02 |
| | | 31 | 44 | Q(-17.03)GVN(+0.98)DNEEGFFSAR | 1.89 | 2.52 |
| | | 33 | 45 | VNDNEEGFFSARG | 1.65 | 2.23 |
| | | 52 | 71 | KREEAPSLRP(+15.99)APPPISGGGY | -1.28 | 2.49 |

| Protein | Accession | Start | End | Sequence | Log2FC | -Log10PV |
|---|---|---|---|---|---|---|
| FIBG | P02679 | 437 | 453 | EHPAETEYDSLYPEDDL | 4.89 | 5.28 |
| FLNA | P21333 | 1048 | 1060 | DGVPVPGSPFPLE | 2.01 | 2.05 |
| FRM4B | Q9Y2L6 | 584 | 601 | DTTTYDDPSDAFTFPGQR | 1.97 | 3.02 |
| FYB1 | O15117 | 44 | 58 | NASPPAGPSNVPKFG | 2.05 | 2.60 |
| FZD8 | Q9H461 | 342 | 368 | GGAPGAGGAGGAGGAAAGAGAAGAGAG | 6.30 | 12.01 |
| GELS | P06396 | 420 | 431 | VPFDAATLHTST | 1.85 | 2.63 |
| | | 420 | 429 | VPFDAATLHT | 1.46 | 1.44 |
| H12 | P16403 | 2 | 21 | S(+42.01)ETAPAAPAAAPPAEKAPVK | 5.49 | 9.86 |
| H14 | P10412 | 2 | 21 | S(+42.01)ETAPAAPAAPAPAEKTPVK | 1.36 | 2.11 |
| H15 | P16401 | 2 | 20 | S(+42.01)ETAPAETATPAPVEKSPA | 1.48 | 1.68 |
| HSPB1 | P04792 | 34 | 42 | GLPRLPEEW | 1.56 | 1.37 |
| | | 177 | 185 | NEITIPVTF | 2.58 | 3.74 |
| | | 190 | 204 | QLGGPEAAKSDETAA | 1.39 | 1.51 |
| IGF2 | P01344 | 93 | 110 | DVSTPPTVLPDNFPRYPV(-0.98) | 3.67 | 4.04 |
| INS | P01308 | 57 | 86 | EAEDLQVGQVELGGGPGAGSLQPLALEGSL | -4.38 | 4.95 |
| | | 57 | 79 | EAEDLQVGQVELGGGPGAGSLQP | 3.89 | 2.76 |
| INSM1 | Q01101 | 135 | 158 | AALLGGGGGGGASGAGGGGTCGGD | 3.71 | 3.31 |
| IRF1 | P10914 | 101 | 109 | KGSSAVRVY | -1.06 | 2.21 |
| ITIH4 | Q14624 | 488 | 500 | GSEMVVAGKLQDR | -1.46 | 1.33 |
| | | 489 | 500 | SEM(+15.99)VVAGKLQDR | 4.08 | 7.30 |
| | | 617 | 625 | N(+0.98)VHSGSTFF | 3.95 | 8.09 |
| | | 635 | 644 | PKPEASFSPR | -2.13 | 2.95 |
| | | 647 | 659 | WNRQAGAAGSRM(+15.99)N | 5.83 | 4.49 |
| | | 660 | 666 | FRPGVLS | 4.21 | 6.97 |
| | | 663 | 683 | GVLSSRQLGLP(+15.99)GPPDVPDHAA | 1.81 | 5.96 |
| | | 669 | 684 | Q(-17.03)LGLPGPPDVPDHAAY | -1.00 | 1.56 |
| | | 673 | 686 | PGPPDVPDHAAYHP | 1.52 | 1.68 |
| | | 673 | 682 | PGPPDVPDHA | -2.58 | 4.06 |
| JUND | P17535 | 153 | 173 | LGAGAAAAAAAAAAGGPSGTA | 1.21 | 1.68 |
| K0513 | O60268 | 238 | 249 | TENVKGFFGGLE | -1.31 | 1.51 |
| KLD10 | Q6PID8 | 17 | 36 | AAGAGGGGSGAGGGSGGSGG | -1.52 | 3.51 |
| LASP1 | Q14847 | 160 | 172 | Q(-17.03)PHHIPTSAPVYQ | 1.63 | 2.13 |
| LEGL | Q3ZCW2 | 2 | 22 | A(+42.01)GSVADSDAVVKLDDGHLN(+0.98)NS | 1.49 | 2.71 |
| | | 7 | 23 | DSDAVVKLDDGHLNNSL | -1.07 | 1.96 |
| LPP | Q93052 | 182 | 212 | PQPAPQAGPIPVAPIGTLKPQPQPVPASYTT | 2.34 | 3.56 |
| LRBA | P50851 | 1568 | 1583 | SENENVSLSEITPAAF | -1.53 | 1.95 |
| LTBP1 | Q14766 | 821 | 834 | GQPQLSPGISTIHL | -1.81 | 2.02 |
| MAGD4 | Q96JG8 | 83 | 90 | QTLVEALQ | 1.41 | 2.67 |
| MGP | P08493 | 21 | 36 | ESHESMESYELNPFIN | 2.16 | 1.70 |
| MMRN1 | Q13201 | 296 | 311 | AESHTAVGRGVAEQQQ | 2.21 | 3.22 |
| MOES | P26038 | 469 | 495 | TPHVAEPAENEQDEQDENGAEASADLR | 2.63 | 4.05 |
| | | 533 | 545 | RDESKKTANDM(+15.99)IH | 3.21 | 4.25 |

| Protein | Accession | Start | End | Sequence | Log2FC | -Log10PV |
|---|---|---|---|---|---|---|
| MYH9 | P35579 | 2 | 11 | A(+42.01)QQAADKYLY | 7.50 | 10.24 |
| | | 1936 | 1955 | RKGAGDGSDEEVDGKADGAE | -1.14 | 1.53 |
| NDUC2 | O95298 | 107 | 115 | TYGEIFEKF | 3.43 | 5.61 |
| OIT3 | Q8WWZ8 | 520 | 545 | GAGGEDSAGLQGQTLTGGPIRIDWED | -4.73 | 8.83 |
| PDLI1 | O00151 | 137 | 173 | ARVITNQYNNPAGLYSSENISNFNNALESKTAASGVE | 1.68 | 2.76 |
| | | 152 | 173 | SSENISNFNN(+0.98)ALESKTAASGVE | 2.93 | 7.30 |
| | | 162 | 173 | ALESKTAASGVE | -3.74 | 7.39 |
| | | 170 | 181 | SGVEANSRPLDH | 1.36 | 1.87 |
| | | 225 | 238 | SEEKGDPNKPSGFR | 2.33 | 2.64 |
| PDLI5 | Q96HC4 | 161 | 174 | ASPSPVAAVTPPLF | 2.66 | 1.6 |
| | | 164 | 174 | SPVAAVTPPLF | 2.33 | 1.68 |
| PDLI7 | Q9NR12 | 100 | 113 | DPPRYTFAPSVSLN | 4.60 | 7.42 |
| PIGR | P01833 | 604 | 639 | LFAEEKAVADTRDQADGSRASVDSGSSEEQGGSSRA | 1.68 | 1.81 |
| PLEC | Q15149 | 3116 | 3125 | SLVPAAELLE | -3.06 | 5.61 |
| PLEK | P08567 | 283 | 292 | AEDPLGAIHL | -7.37 | 16.16 |
| PRRC1 | Q96M27 | 164 | 180 | TGLLPTPITQQASLTSL | 2.67 | 5.61 |
| PTN12 | Q05209 | 568 | 591 | LTPSPTTQVETPDLVDHDNTSPLF | -1.09 | 1.42 |
| RTN4 | Q9NQC3 | 97 | 124 | APPVAPERQPSWDPSPVSSTVPAPSPLS(+21.98) | 3.08 | 3.20 |
| S10A8 | P05109 | 18 | 26 | KYSLIKGNF | 2.69 | 2.87 |
| SAA2 | P0DJI9 | 109 | 122 | DPNHFRPAGLPEKY | -3.43 | 11.83 |
| SCAM2 | O15127 | 31 | 53 | QGGLAEFNPFSETNAATTVPVTQ | -1.00 | 2.25 |
| SKAP2 | O75563 | 253 | 277 | SQPIDDEIYEELPEEEEDSAPVKVE | -1.16 | 2.66 |
| SKOR1 | P84550 | 292 | 319 | GGSGGQGKGGAGGGGGGGPGCGAEM(+15.99)APG | 3.55 | 3.40 |
| SRC8 | Q14247 | 130 | 141 | DRVDQSAVGFEY | -1.82 | 4.11 |
| SYUA | P37840 | 32 | 47 | KTKEGVLYVGSKTKEG | 1.46 | 4.78 |
| TAGL2 | P37802 | 2 | 12 | A(+42.01)NRGPAYGLSR | 3.98 | 4.89 |
| | | 2 | 11 | A(+42.01)NRGPAYGLS(-18.01) | 2.82 | 4.60 |
| | | 180 | 199 | TNRGASQAGM(+15.99)TGYGM(+15.99)PRQIL | -3.09 | 3.20 |
| | | 180 | 192 | TNRGASQAGM(+15.99)TGY | 4.13 | 3.50 |
| | | 181 | 197 | NRGASQAGMTGYGMPRQ | 4.39 | 7.76 |
| TBA1B | P68363 | 29 | 57 | GIQPDGQMPSDKTIGGGDDSFNTFFSETG | -1.58 | 3.00 |
| | | 37 | 57 | PSDKTIGGGDDSFNTFFSETG | -3.23 | 3.59 |
| | | 66 | 79 | VFVDLEPTVIDEVR | 1.82 | 2.16 |
| | | 169 | 178 | FSIYPAPQVS | 1.83 | 2.86 |
| | | 230 | 240 | LISQIVSSITA | 1.71 | 5.16 |
| | | 276 | 285 | ISAEKAYHEQ | 1.10 | 1.31 |
| | | 283 | 293 | HEQLSVAEITN | -1.16 | 1.77 |
| | | 381 | 392 | TTAIAEAWARLD | 3.15 | 5.88 |
| | | 382 | 392 | TAIAEAWARLD | -2.68 | 9.73 |
| | | 430 | 450 | KDYEEVGVDS(-18.01)VEGEGEEEGEE | 4.68 | 9.73 |
| TBA1C | Q9BQE3 | 379 | 392 | SNTTAVAEAWARLD | -1.64 | 2 |
| | | 430 | 449 | KDYEEVGADSADGEDEGEEY | 1.26 | 2.24 |
| TBA4A | P68366 | 38 | 52 | SDKTIGGGDDSFTTF | -1.65 | 1.66 |

| Protein | Accession | Start | End | Sequence | Log2FC | -Log10PV |
|---|---|---|---|---|---|---|
| | | 430 | 446 | KDYEEVGIDSYEDEDEG | 1.37 | 2.22 |
| | | 437 | 448 | IDSYEDEDEGEE | 1.00 | 1.51 |
| TBA8 | Q9NY65 | 41 | 57 | SKINDDDSFTTFFSETG | -1.18 | 2.43 |
| | | 43 | 57 | INDDDSFTTFFSETG | 2.97 | 5.21 |
| | | 43 | 52 | INDDDSFTTF | -1.28 | 2.67 |
| | | 276 | 285 | ISAEKAYHEQ | 1.10 | 1.31 |
| TBB1 | Q9H4B7 | 107 | 118 | TEGAELIENVLE | 1.90 | 1.89 |
| | | 168 | 192 | SVMPSPKVSDTVVEPYNAVLSIHQL | -1.26 | 1.45 |
| | | 168 | 187 | SVMPSPKVSDTVVEPYNAVL | -1.48 | 1.55 |
| | | 218 | 232 | TTPTYGDLNHLVSLT | 2.82 | 2.4 |
| | | 403 | 419 | MDINEFGEAENNIHDLV | 1.00 | 2.24 |
| TBB5 | P07437 | 14 | 25 | N(+0.98)QIGAKFWEVIS | -1.30 | 2.64 |
| | | 16 | 25 | IGAKFWEVIS | 2.47 | 3.29 |
| | | 106 | 121 | YTEGAELVDSVLDVVR | 3.44 | 5.59 |
| | | 123 | 136 | EAES(-18.01)CD(+15.99)CLQGFQLT | 2.03 | 2.43 |
| | | 224 | 238 | DLNHLVSATMSGVTT | -1.40 | 2.90 |
| | | 420 | 443 | SEYQQYQDATAEEEED(-18.01)FGEEAEEE | 3.61 | 6.83 |
| | | 420 | 441 | SEYQQYQDATAEEEED(-18.01)FGEEAE | 3.40 | 7.66 |
| | | 423 | 444 | Q(-17.03)QYQDATAEEEEDFGEEAEEEA | 1.18 | 1.40 |
| TCRG1 | O14776 | 255 | 271 | AVGASTPTTSSPAPAVS | 2.41 | 2.53 |
| TLN1 | Q9Y490 | 409 | 427 | FGLEGDEESTMLEDSVSPK | -1.48 | 2.76 |
| | | 436 | 446 | YNRVGKVEHGS(-18.01) | -2.30 | 3.61 |
| | | 873 | 891 | AAKGAAAHPDSEEQQQRLR | 1.08 | 1.92 |
| | | 1531 | 1541 | EVANSTANLVK | -1.24 | 2.11 |
| | | 1560 | 1575 | AATAPLLEAVDNLSAF | 1.55 | 1.86 |
| | | 1569 | 1581 | VDNLSAFASNPEF | 4.64 | 7.87 |
| | | 2512 | 2519 | ERELEEAR | 3.90 | 1.63 |
| TMOD3 | Q9NYL9 | 170 | 177 | GEKILPVF | -1.73 | 3.01 |
| TRML1 | Q86YW5 | 218 | 235 | DSGPAAELPLDVPHIRLD | 4.89 | 7.51 |
| | | 234 | 251 | LDSPPSFDNTTYTSLPLD | 2.18 | 5.53 |
| | | 246 | 274 | TSLPLDSPSGKPSLPAPSSLPPLPPKVLV | 3.83 | 6.42 |
| | | 247 | 273 | SLPLDSPSGKPSLPAPSSLPPLPPKVL | 5.63 | 11.83 |
| TYB10 | P63313 | 2 | 18 | ADK(+42.02)PDMGEIASFDKAKL | -4.65 | 5.57 |
| TYB4 | P62328 | 2 | 27 | SDK(+42.01)PDMAEIEKFDKSKLKKT(-18.01)ETQEKN | 7.13 | 11.01 |
| | | 2 | 25 | S(+42.01)DKPDM(+15.99)AEIEKFDKSKLKKTETQE | 7.66 | 10.06 |
| | | 2 | 23 | S(+42.01)DKPDMAEIEKFDKSKLKKTET | 2.72 | 2.43 |
| | | 2 | 22 | SDK(+42.02)PDMAEIEKFDKSKLKKTE | 3.79 | 2.44 |
| | | 2 | 22 | S(+42.01)DKPDMAEIEKFDKSKLKKTE | 8.83 | 8.36 |
| | | 26 | 38 | KNPLPSKET(-18.01)IEQE | 2.83 | 5.22 |
| UBQL1 | Q9UMX0 | 293 | 308 | GNPFASLVSNTSSGEG | 3.65 | 7.25 |
| URP2 | Q86UX7 | 338 | 352 | APTDVLDSLTTIPEL | -1.21 | 1.54 |
| | | 341 | 352 | DVLDSLTTIPEL | -2.95 | 6.95 |
| VASP | P50552 | 128 | 141 | SVPNGPSPEEVEQQ | 2.55 | 2.81 |

| Protein | Accession | Start | End | Sequence | Log2FC | -Log10PV |
|---|---|---|---|---|---|---|
| VIME | P08670 | 30 | 38 | YVTTSTRTY | -1.86 | 1.44 |
| VTNC | P04004 | 94 | 115 | GGPSLTSDLQAQSKGNPEQTPV | 2.80 | 4.85 |
| | | 97 | 115 | SLTSDLQAQSKGNPEQTPV | 6.45 | 9.82 |
| | | 108 | 128 | GNPEQTPVLKPEEEAPAPEVG | 1.84 | 2.69 |
| | | 117 | 135 | KPEEEAPAPEVGASKPEGI | 4.84 | 8.94 |
| | | 119 | 135 | EEEAPAPEVGASKPEGI | 6.99 | 7.23 |
| WDR44 | Q5JSH3 | 63 | 72 | IIESIIEESQ | 2.21 | 2.54 |
| ZYX | Q15942 | 247 | 279 | LANTQPRGPPASSPAPAPKFSPVTPKFTPVASK | 1.01 | 1.49 |
| | | 345 | 360 | PGAPGPLTLKEVEELE | -3.39 | 8.33 |

Log2FC and -Log10PV are in comparison to values from patients that did not progress to >F2 fibrosis. Start and end of the peptide are based on the FASTA protein sequence associated with the accession number. Mass addition (e.g., +42.02) and mass loss (e.g., -18.01) denotes chemical modification of the amino acid, corresponding to the following:

| | |
|---|---|
| -18.01 | dehydration |
| -17.03 | pyroglutamic acid |
| -0.98 | amidation |
| +0.98 | deamidation |
| +15.99 | oxidation |
| +18.01 | hydration |
| +21.98 | sodiation |
| +27.99 | formylation |
| +28.03 | dimethylation |
| +42.01 | acetylation |
| +42.02 | guanidiation |
| +43.01 | carbamylation |
| +79.97 | phosphorylation |
| +156.12 | 4-hydroxynonenal |
| +162.05 | hexose |

**Supplemental Table 4-GO terms for Cellular Component (GO:0005575) of peptides changed in post-LT fibrosis**

| Term ID # | Term Description | Count | Enrichment | -Log10PV |
|---|---|---|---|---|
| **Increased in post LT fibrosis** | | | | |
| GO:0015629 | actin cytoskeleton | 20 | 0.99 | 11.06 |
| GO:0034774 | secretory granule lumen | 18 | 1.07 | 11.06 |
| GO:0005856 | cytoskeleton | 37 | 0.58 | 10.75 |
| GO:0099503 | secretory vesicle | 26 | 0.76 | 10.75 |
| GO:0030141 | secretory granule | 24 | 0.79 | 10.36 |
| GO:0005938 | cell cortex | 14 | 1.11 | 9.42 |
| GO:0005747 | mitochondrial respiratory chain complex I | 8 | 1.56 | 8.21 |
| GO:0043229 | intracellular organelle | 85 | 0.17 | 8.21 |
| GO:0030863 | cortical cytoskeleton | 9 | 1.39 | 8.15 |
| GO:0031410 | cytoplasmic vesicle | 34 | 0.51 | 8.15 |
| GO:0005737 | cytoplasm | 81 | 0.18 | 7.88 |
| GO:0099568 | cytoplasmic region | 15 | 0.89 | 7.72 |
| GO:0005576 | extracellular region | 34 | 0.46 | 7.01 |
| GO:0005622 | intracellular | 89 | 0.12 | 6.7 |
| GO:0005788 | endoplasmic reticulum lumen | 12 | 0.93 | 6.45 |
| GO:0043232 | intracellular non-membrane-bounded organelle | 43 | 0.35 | 6.45 |
| GO:0099081 | supramolecular polymer | 19 | 0.66 | 6.38 |
| GO:0098644 | complex of collagen trimers | 5 | 1.74 | 6.03 |
| GO:0032991 | protein-containing complex | 46 | 0.31 | 5.7 |
| GO:0012505 | endomembrane system | 43 | 0.32 | 5.55 |
| GO:0005581 | collagen trimer | 7 | 1.22 | 5.49 |
| GO:0030864 | cortical actin cytoskeleton | 6 | 1.38 | 5.48 |
| GO:0031091 | platelet alpha granule | 7 | 1.21 | 5.42 |
| GO:0002102 | podosome | 5 | 1.54 | 5.26 |
| GO:0099512 | supramolecular fiber | 17 | 0.61 | 5.14 |
| GO:0031093 | platelet alpha granule lumen | 6 | 1.27 | 4.94 |
| GO:0005925 | focal adhesion | 7 | 1.04 | 4.4 |
| GO:0005924 | cell-substrate adherens junction | 7 | 1.03 | 4.36 |
| GO:0030054 | cell junction | 17 | 0.55 | 4.36 |
| GO:0001726 | ruffle | 7 | 0.95 | 3.89 |
| GO:0005577 | fibrinogen complex | 3 | 1.9 | 3.89 |
| GO:0042641 | actomyosin | 5 | 1.22 | 3.89 |
| GO:0005615 | extracellular space | 17 | 0.5 | 3.74 |
| GO:0005911 | cell-cell junction | 10 | 0.72 | 3.74 |
| GO:0005912 | adherens junction | 8 | 0.82 | 3.66 |
| GO:0005583 | fibrillar collagen trimer | 3 | 1.76 | 3.62 |
| GO:0005623 | cell | 90 | 0.07 | 3.52 |
| GO:0062023 | collagen-containing extracellular matrix | 6 | 0.94 | 3.33 |
| GO:0070013 | intracellular organelle lumen | 42 | 0.23 | 3.33 |
| GO:0043227 | membrane-bounded organelle | 71 | 0.12 | 3.17 |
| GO:0001725 | stress fiber | 4 | 1.23 | 3.16 |
| GO:0005584 | collagen type I trimer | 2 | 2.32 | 3.14 |
| GO:0071944 | cell periphery | 41 | 0.22 | 2.85 |
| GO:0101002 | ficolin-1-rich granule | 6 | 0.83 | 2.8 |
| GO:0030478 | actin cap | 2 | 2.02 | 2.77 |
| GO:0005743 | mitochondrial inner membrane | 9 | 0.62 | 2.74 |
| GO:0005886 | plasma membrane | 40 | 0.21 | 2.72 |
| GO:0031966 | mitochondrial membrane | 11 | 0.53 | 2.7 |
| GO:0031252 | cell leading edge | 8 | 0.66 | 2.68 |
| GO:0031012 | extracellular matrix | 7 | 0.72 | 2.66 |
| GO:0005874 | microtubule | 8 | 0.64 | 2.59 |
| GO:0005587 | collagen type IV trimer | 2 | 1.85 | 2.55 |
| GO:0034363 | intermediate-density lipoprotein particle | 2 | 1.85 | 2.55 |
| GO:0042995 | cell projection | 20 | 0.33 | 2.43 |

| Term ID # | Term Description | Count | Enrichment | -Log10PV |
|---|---|---|---|---|
| GO:0042582 | azurophil granule | 5 | 0.83 | 2.39 |
| GO:0034366 | spherical high-density lipoprotein particle | 2 | 1.67 | 2.32 |
| GO:0098794 | postsynapse | 8 | 0.59 | 2.32 |
| GO:0120025 | plasma membrane bounded cell projection | 19 | 0.32 | 2.24 |
| GO:0001931 | uropod | 2 | 1.51 | 2.06 |
| GO:0042627 | chylomicron | 2 | 1.51 | 2.06 |
| GO:0045202 | synapse | 11 | 0.44 | 2.03 |
| GO:0031967 | organelle envelope | 13 | 0.38 | 1.93 |
| GO:0014069 | postsynaptic density | 5 | 0.71 | 1.93 |
| GO:1904813 | ficolin-1-rich granule lumen | 4 | 0.83 | 1.92 |
| GO:0031143 | pseudopodium | 2 | 1.39 | 1.9 |
| GO:0005905 | clathrin-coated pit | 3 | 1 | 1.88 |
| GO:0099513 | polymeric cytoskeletal fiber | 9 | 0.47 | 1.88 |
| GO:0030016 | myofibril | 5 | 0.69 | 1.87 |
| GO:0030425 | dendrite | 8 | 0.5 | 1.87 |
| GO:0005783 | endoplasmic reticulum | 17 | 0.3 | 1.82 |
| GO:0016020 | membrane | 53 | 0.12 | 1.81 |
| GO:0034361 | very-low-density lipoprotein particle | 2 | 1.32 | 1.81 |
| GO:0030659 | cytoplasmic vesicle membrane | 9 | 0.42 | 1.61 |
| GO:0036477 | somatodendritic compartment | 9 | 0.41 | 1.59 |
| GO:0070820 | tertiary granule | 4 | 0.71 | 1.59 |
| GO:0031970 | organelle envelope lumen | 3 | 0.86 | 1.57 |
| GO:0005604 | basement membrane | 3 | 0.84 | 1.52 |
| GO:0035578 | azurophil granule lumen | 3 | 0.84 | 1.51 |
| GO:0098796 | membrane protein complex | 11 | 0.34 | 1.49 |
| GO:0030027 | lamellipodium | 4 | 0.66 | 1.44 |
| GO:0005720 | nuclear heterochromatin | 2 | 1.08 | 1.42 |
| GO:0032993 | protein-DNA complex | 4 | 0.65 | 1.42 |
| GO:0000786 | nucleosome | 3 | 0.77 | 1.37 |
| GO:0048471 | perinuclear region of cytoplasm | 8 | 0.4 | 1.37 |
| GO:0098805 | whole membrane | 14 | 0.28 | 1.37 |
| GO:0005773 | vacuole | 8 | 0.39 | 1.34 |
| GO:0015630 | microtubule cytoskeleton | 11 | 0.32 | 1.33 |
| **Decreased in post LT fibrosis** | | | | |
| GO:0005856 | cytoskeleton | 22 | 0.63 | 6.58 |
| GO:0099080 | supramolecular complex | 15 | 0.83 | 6.48 |
| GO:0015629 | actin cytoskeleton | 11 | 1.01 | 6.1 |
| GO:0099512 | supramolecular fiber | 14 | 0.81 | 5.88 |
| GO:0031093 | platelet alpha granule lumen | 5 | 1.47 | 4.35 |
| GO:0002102 | podosome | 4 | 1.73 | 4.26 |
| GO:0005938 | cell cortex | 7 | 1.08 | 4.26 |
| GO:0034774 | secretory granule lumen | 8 | 1 | 4.26 |
| GO:0005577 | fibrinogen complex | 3 | 2.18 | 4.26 |
| GO:0005576 | extracellular region | 19 | 0.48 | 4.09 |
| GO:0005737 | cytoplasm | 43 | 0.18 | 4.09 |
| GO:0043232 | intracellular non-membrane-bounded organelle | 24 | 0.38 | 3.89 |
| GO:0030141 | secretory granule | 10 | 0.68 | 3.31 |
| GO:0030863 | cortical cytoskeleton | 4 | 1.32 | 3.17 |
| GO:0099568 | cytoplasmic region | 7 | 0.84 | 3.08 |
| GO:0005622 | intracellular | 46 | 0.11 | 2.59 |
| GO:0031410 | cytoplasmic vesicle | 15 | 0.43 | 2.55 |
| GO:0043292 | contractile fiber | 5 | 0.94 | 2.54 |
| GO:0031252 | cell leading edge | 6 | 0.81 | 2.48 |
| GO:0005874 | microtubule | 6 | 0.79 | 2.43 |
| GO:0005924 | cell-substrate adherens junction | 4 | 1.07 | 2.43 |
| GO:0005925 | focal adhesion | 4 | 1.08 | 2.43 |
| GO:0042641 | actomyosin | 3 | 1.28 | 2.32 |
| GO:0001726 | ruffle | 4 | 0.99 | 2.2 |

| Term ID # | Term Description | Count | Enrichment | -Log10PV |
|---|---|---|---|---|
| GO:0043229 | intracellular organelle | 41 | 0.13 | 2.12 |
| GO:0099513 | polymeric cytoskeletal fiber | 7 | 0.64 | 2.09 |
| GO:0098644 | complex of collagen trimers | 2 | 1.62 | 2.06 |
| GO:0005615 | extracellular space | 9 | 0.5 | 1.9 |
| GO:0030016 | myofibril | 4 | 0.87 | 1.84 |
| GO:0034364 | high-density lipoprotein particle | 2 | 1.46 | 1.79 |
| GO:0005912 | adherens junction | 4 | 0.8 | 1.64 |
| GO:0032991 | protein-containing complex | 21 | 0.24 | 1.64 |
| GO:0031012 | extracellular matrix | 4 | 0.75 | 1.51 |
| GO:0062023 | collagen-containing extracellular matrix | 3 | 0.92 | 1.5 |
| GO:0015630 | microtubule cytoskeleton | 8 | 0.46 | 1.48 |
| GO:0005788 | endoplasmic reticulum lumen | 4 | 0.73 | 1.46 |
| GO:0001725 | stress fiber | 2 | 1.2 | 1.43 |
| GO:0030864 | cortical actin cytoskeleton | 2 | 1.18 | 1.4 |

**Supplemental Table 5-GO terms for Biological Process (GO:0008150) of peptides changed in post-LT fibrosis**

| Term ID # | Term Description | Count | Enrichment | -Log10PV |
|---|---|---|---|---|
| **Increased in post LT fibrosis** | | | | |
| GO:0045055 | regulated exocytosis | 25 | 0.88 | 11.30 |
| GO:0001775 | cell activation | 27 | 0.74 | 9.96 |
| GO:0002576 | platelet degranulation | 13 | 1.33 | 9.96 |
| GO:0016192 | vesicle-mediated transport | 34 | 0.62 | 9.96 |
| GO:0016043 | cellular component organization | 56 | 0.36 | 8.53 |
| GO:0022607 | cellular component assembly | 37 | 0.52 | 8.47 |
| GO:0007010 | cytoskeleton organization | 23 | 0.71 | 7.62 |
| GO:0006120 | mitochondrial electron transport, NADH to ubiquinone | 8 | 1.58 | 7.62 |
| GO:0065003 | protein-containing complex assembly | 28 | 0.59 | 7.32 |
| GO:0042775 | mitochondrial ATP synthesis coupled electron transport | 9 | 1.39 | 7.30 |
| GO:0043933 | protein-containing complex subunit organization | 30 | 0.55 | 7.23 |
| GO:0006996 | organelle organization | 40 | 0.43 | 7.01 |
| GO:0030036 | actin cytoskeleton organization | 15 | 0.88 | 6.66 |
| GO:0032981 | mitochondrial respiratory chain complex I assembly | 8 | 1.41 | 6.64 |
| GO:0030168 | platelet activation | 9 | 1.2 | 5.97 |
| GO:0034622 | cellular protein-containing complex assembly | 19 | 0.68 | 5.85 |
| GO:1900026 | positive regulation of substrate adhesion-dependent cell spreading | 6 | 1.6 | 5.67 |
| GO:0050776 | regulation of immune response | 19 | 0.66 | 5.58 |
| GO:0097435 | supramolecular fiber organization | 13 | 0.85 | 5.43 |
| GO:0048584 | positive regulation of response to stimulus | 29 | 0.47 | 5.39 |
| GO:0051130 | positive regulation of cellular component organization | 21 | 0.59 | 5.27 |
| GO:0002376 | immune system process | 31 | 0.44 | 5.21 |
| GO:0002682 | regulation of immune system process | 23 | 0.54 | 5.07 |
| GO:0051234 | establishment of localization | 43 | 0.33 | 5.00 |
| GO:0007160 | cell-matrix adhesion | 8 | 1.15 | 4.98 |
| GO:0006897 | endocytosis | 14 | 0.76 | 4.97 |
| GO:0006810 | transport | 42 | 0.33 | 4.92 |
| GO:0045321 | leukocyte activation | 18 | 0.63 | 4.87 |
| GO:0051258 | protein polymerization | 7 | 1.25 | 4.87 |
| GO:0042060 | wound healing | 13 | 0.77 | 4.70 |
| GO:0009611 | response to wounding | 14 | 0.73 | 4.68 |
| GO:0010770 | positive regulation of cell morphogenesis involved in differentiation | 8 | 1.09 | 4.65 |
| GO:0051128 | regulation of cellular component organization | 29 | 0.42 | 4.51 |
| GO:0043312 | neutrophil degranulation | 13 | 0.75 | 4.50 |
| GO:0007015 | actin filament organization | 9 | 0.98 | 4.49 |
| GO:0002274 | myeloid leukocyte activation | 14 | 0.71 | 4.49 |
| GO:0010769 | regulation of cell morphogenesis involved in differentiation | 10 | 0.9 | 4.49 |
| GO:0043062 | extracellular structure organization | 11 | 0.83 | 4.43 |
| GO:0051179 | localization | 47 | 0.28 | 4.38 |
| GO:0010811 | positive regulation of cell-substrate adhesion | 7 | 1.13 | 4.28 |
| GO:0098657 | import into cell | 14 | 0.68 | 4.28 |
| GO:0007596 | blood coagulation | 10 | 0.86 | 4.26 |
| GO:0009205 | purine ribonucleoside triphosphate metabolic process | 9 | 0.93 | 4.26 |
| GO:0022604 | regulation of cell morphogenesis | 12 | 0.76 | 4.25 |
| GO:0010647 | positive regulation of cell communication | 23 | 0.47 | 4.19 |
| GO:0009167 | purine ribonucleoside monophosphate metabolic process | 9 | 0.92 | 4.18 |
| GO:0002443 | leukocyte mediated immunity | 14 | 0.67 | 4.18 |
| GO:0023056 | positive regulation of signaling | 23 | 0.47 | 4.18 |
| GO:0072376 | protein activation cascade | 6 | 1.23 | 4.16 |

| Term ID # | Term Description | Count | Enrichment | -Log10PV |
|---|---|---|---|---|
| GO:2001233 | regulation of apoptotic signaling pathway | 11 | 0.78 | 4.04 |
| GO:0022603 | regulation of anatomical structure morphogenesis | 17 | 0.57 | 4.04 |
| GO:0006955 | immune response | 22 | 0.47 | 4.00 |
| GO:0010810 | regulation of cell-substrate adhesion | 8 | 0.95 | 3.89 |
| GO:0031347 | regulation of defense response | 14 | 0.64 | 3.89 |
| GO:0050793 | regulation of developmental process | 28 | 0.39 | 3.85 |
| GO:0034114 | regulation of heterotypic cell-cell adhesion | 4 | 1.6 | 3.80 |
| GO:0050727 | regulation of inflammatory response | 10 | 0.79 | 3.80 |
| GO:0009967 | positive regulation of signal transduction | 21 | 0.47 | 3.77 |
| GO:0002252 | immune effector process | 16 | 0.56 | 3.66 |
| GO:0048583 | regulation of response to stimulus | 37 | 0.3 | 3.66 |
| GO:0006898 | receptor-mediated endocytosis | 8 | 0.91 | 3.64 |
| GO:2000257 | regulation of protein activation cascade | 5 | 1.29 | 3.64 |
| GO:0051235 | maintenance of location | 7 | 0.99 | 3.60 |
| GO:0032101 | regulation of response to external stimulus | 16 | 0.55 | 3.52 |
| GO:0010638 | positive regulation of organelle organization | 12 | 0.66 | 3.46 |
| GO:0030198 | extracellular matrix organization | 9 | 0.81 | 3.46 |
| GO:0060627 | regulation of vesicle-mediated transport | 11 | 0.68 | 3.30 |
| GO:0050878 | regulation of body fluid levels | 11 | 0.68 | 3.28 |
| GO:0006911 | phagocytosis, engulfment | 4 | 1.43 | 3.28 |
| GO:0038063 | collagen-activated tyrosine kinase receptor signaling pathway | 3 | 1.85 | 3.28 |
| GO:0051050 | positive regulation of transport | 15 | 0.55 | 3.28 |
| GO:0070613 | regulation of protein processing | 6 | 1.04 | 3.24 |
| GO:0051246 | regulation of protein metabolic process | 28 | 0.34 | 3.14 |
| GO:0032879 | regulation of localization | 27 | 0.35 | 3.12 |
| GO:0031639 | plasminogen activation | 3 | 1.76 | 3.09 |
| GO:0007005 | mitochondrion organization | 10 | 0.7 | 3.06 |
| GO:0032970 | regulation of actin filament-based process | 9 | 0.75 | 3.06 |
| GO:0080134 | regulation of response to stress | 18 | 0.46 | 3.06 |
| GO:0007155 | cell adhesion | 14 | 0.54 | 3.00 |
| GO:1903035 | negative regulation of response to wounding | 5 | 1.12 | 3.00 |
| GO:0034116 | positive regulation of heterotypic cell-cell adhesion | 3 | 1.69 | 2.96 |
| GO:0034329 | cell junction assembly | 6 | 0.97 | 2.96 |
| GO:0051247 | positive regulation of protein metabolic process | 20 | 0.42 | 2.96 |
| GO:1902533 | positive regulation of intracellular signal transduction | 15 | 0.52 | 2.96 |
| GO:0007229 | integrin-mediated signaling pathway | 5 | 1.1 | 2.89 |
| GO:0008154 | actin polymerization or depolymerization | 4 | 1.29 | 2.85 |
| GO:0032270 | positive regulation of cellular protein metabolic process | 19 | 0.43 | 2.85 |
| GO:0071560 | cellular response to transforming growth factor beta stimulus | 6 | 0.96 | 2.85 |
| GO:0001932 | regulation of protein phosphorylation | 18 | 0.44 | 2.82 |
| GO:0090066 | regulation of anatomical structure size | 10 | 0.66 | 2.80 |
| GO:0019220 | regulation of phosphate metabolic process | 20 | 0.4 | 2.77 |
| GO:0022008 | neurogenesis | 19 | 0.42 | 2.77 |
| GO:0002673 | regulation of acute inflammatory response | 5 | 1.06 | 2.74 |
| GO:0032880 | regulation of protein localization | 14 | 0.51 | 2.74 |
| GO:2001234 | negative regulation of apoptotic signaling pathway | 7 | 0.83 | 2.74 |
| GO:0030195 | negative regulation of blood coagulation | 4 | 1.24 | 2.72 |
| GO:0045597 | positive regulation of cell differentiation | 14 | 0.51 | 2.72 |
| GO:0033043 | regulation of organelle organization | 16 | 0.46 | 2.70 |
| GO:0034097 | response to cytokine | 15 | 0.48 | 2.70 |
| GO:0045185 | maintenance of protein location | 5 | 1.04 | 2.70 |
| GO:0051094 | positive regulation of developmental process | 17 | 0.44 | 2.70 |
| GO:0090277 | positive regulation of peptide hormone secretion | 5 | 1.04 | 2.70 |
| GO:0002253 | activation of immune response | 9 | 0.68 | 2.68 |
| GO:0045595 | regulation of cell differentiation | 20 | 0.39 | 2.68 |

| Term ID # | Term Description | Count | Enrichment | -Log10PV |
|---|---|---|---|---|
| GO:0048518 | positive regulation of biological process | 43 | 0.22 | 2.66 |
| GO:0051621 | regulation of norepinephrine uptake | 2 | 2.32 | 2.66 |
| GO:1903923 | positive regulation of protein processing in phagocytic vesicle | 2 | 2.32 | 2.66 |
| GO:0009653 | anatomical structure morphogenesis | 22 | 0.37 | 2.64 |
| GO:0030449 | regulation of complement activation | 4 | 1.21 | 2.64 |
| GO:0065008 | regulation of biological quality | 32 | 0.28 | 2.62 |
| GO:0110053 | regulation of actin filament organization | 7 | 0.8 | 2.62 |
| GO:1902903 | regulation of supramolecular fiber organization | 8 | 0.73 | 2.62 |
| GO:2001237 | negative regulation of extrinsic apoptotic signaling pathway | 5 | 1 | 2.59 |
| GO:0043410 | positive regulation of MAPK cascade | 10 | 0.61 | 2.55 |
| GO:0051239 | regulation of multicellular organismal process | 27 | 0.31 | 2.55 |
| GO:0031399 | regulation of protein modification process | 20 | 0.38 | 2.54 |
| GO:0032268 | regulation of cellular protein metabolic process | 25 | 0.33 | 2.54 |
| GO:0042730 | fibrinolysis | 3 | 1.48 | 2.54 |
| GO:0031532 | actin cytoskeleton reorganization | 4 | 1.17 | 2.52 |
| GO:0060284 | regulation of cell development | 13 | 0.51 | 2.52 |
| GO:0009150 | purine ribonucleotide metabolic process | 9 | 0.65 | 2.49 |
| GO:0010903 | negative regulation of very-low-density lipoprotein particle remodeling | 2 | 2.15 | 2.49 |
| GO:0043254 | regulation of protein complex assembly | 9 | 0.65 | 2.49 |
| GO:0030162 | regulation of proteolysis | 12 | 0.53 | 2.48 |
| GO:0048468 | cell development | 18 | 0.4 | 2.48 |
| GO:0010646 | regulation of cell communication | 30 | 0.28 | 2.44 |
| GO:0030334 | regulation of cell migration | 12 | 0.53 | 2.44 |
| GO:0044087 | regulation of cellular component biogenesis | 13 | 0.5 | 2.44 |
| GO:1902905 | positive regulation of supramolecular fiber organization | 6 | 0.85 | 2.44 |
| GO:0048522 | positive regulation of cellular process | 39 | 0.22 | 2.41 |
| GO:0002793 | positive regulation of peptide secretion | 7 | 0.75 | 2.40 |
| GO:0040012 | regulation of locomotion | 13 | 0.49 | 2.40 |
| GO:0007165 | signal transduction | 38 | 0.23 | 2.39 |
| GO:0009966 | regulation of signal transduction | 28 | 0.29 | 2.39 |
| GO:0010954 | positive regulation of protein processing | 3 | 1.4 | 2.39 |
| GO:0023051 | regulation of signaling | 30 | 0.27 | 2.39 |
| GO:0032507 | maintenance of protein location in cell | 4 | 1.12 | 2.39 |
| GO:0048869 | cellular developmental process | 31 | 0.27 | 2.39 |
| GO:0070527 | platelet aggregation | 3 | 1.4 | 2.39 |
| GO:0002791 | regulation of peptide secretion | 9 | 0.62 | 2.36 |
| GO:0030335 | positive regulation of cell migration | 9 | 0.62 | 2.36 |
| GO:0040011 | locomotion | 15 | 0.44 | 2.36 |
| GO:0050896 | response to stimulus | 54 | 0.16 | 2.36 |
| GO:0031401 | positive regulation of protein modification process | 15 | 0.44 | 2.35 |
| GO:0072378 | blood coagulation, fibrin clot formation | 3 | 1.37 | 2.32 |
| GO:0045087 | innate immune response | 11 | 0.53 | 2.30 |
| GO:0009605 | response to external stimulus | 22 | 0.33 | 2.29 |
| GO:0035025 | positive regulation of Rho protein signal transduction | 3 | 1.35 | 2.29 |
| GO:0048870 | cell motility | 13 | 0.48 | 2.29 |
| GO:2000352 | negative regulation of endothelial cell apoptotic process | 3 | 1.35 | 2.29 |
| GO:0045807 | positive regulation of endocytosis | 5 | 0.91 | 2.28 |
| GO:0062013 | positive regulation of small molecule metabolic process | 5 | 0.91 | 2.28 |
| GO:0045785 | positive regulation of cell adhesion | 8 | 0.65 | 2.26 |
| GO:0045937 | positive regulation of phosphate metabolic process | 14 | 0.45 | 2.26 |
| GO:0051495 | positive regulation of cytoskeleton organization | 6 | 0.79 | 2.26 |
| GO:0016477 | cell migration | 12 | 0.49 | 2.24 |

| Term ID # | Term Description | Count | Enrichment | -Log10PV |
|---|---|---|---|---|
| GO:0051493 | regulation of cytoskeleton organization | 9 | 0.6 | 2.24 |
| GO:1902175 | regulation of oxidative stress-induced intrinsic apoptotic signaling pathway | 3 | 1.32 | 2.24 |
| GO:0007167 | enzyme linked receptor protein signaling pathway | 11 | 0.52 | 2.24 |
| GO:0010033 | response to organic substance | 26 | 0.29 | 2.23 |
| GO:0001934 | positive regulation of protein phosphorylation | 13 | 0.46 | 2.22 |
| GO:0007166 | cell surface receptor signaling pathway | 22 | 0.32 | 2.21 |
| GO:0070887 | cellular response to chemical stimulus | 25 | 0.29 | 2.19 |
| GO:1902531 | regulation of intracellular signal transduction | 19 | 0.36 | 2.19 |
| GO:0042989 | sequestering of actin monomers | 2 | 1.85 | 2.18 |
| GO:0043152 | induction of bacterial agglutination | 2 | 1.85 | 2.18 |
| GO:0045921 | positive regulation of exocytosis | 4 | 1.03 | 2.18 |
| GO:0051047 | positive regulation of secretion | 8 | 0.63 | 2.18 |
| GO:0071310 | cellular response to organic substance | 22 | 0.32 | 2.18 |
| GO:0071800 | podosome assembly | 2 | 1.85 | 2.18 |
| GO:0098532 | histone H3-K27 trimethylation | 2 | 1.85 | 2.18 |
| GO:1904035 | regulation of epithelial cell apoptotic process | 4 | 1.03 | 2.18 |
| GO:0006928 | movement of cell or subcellular component | 16 | 0.4 | 2.17 |
| GO:0007017 | microtubule-based process | 10 | 0.54 | 2.17 |
| GO:0032989 | cellular component morphogenesis | 11 | 0.51 | 2.17 |
| GO:0045907 | positive regulation of vasoconstriction | 3 | 1.28 | 2.17 |
| GO:0072358 | cardiovascular system development | 9 | 0.58 | 2.17 |
| GO:0032103 | positive regulation of response to external stimulus | 9 | 0.58 | 2.16 |
| GO:0090207 | regulation of triglyceride metabolic process | 3 | 1.27 | 2.14 |
| GO:0061024 | membrane organization | 11 | 0.5 | 2.13 |
| GO:0071495 | cellular response to endogenous stimulus | 14 | 0.43 | 2.12 |
| GO:0032102 | negative regulation of response to external stimulus | 7 | 0.68 | 2.11 |
| GO:0032489 | regulation of Cdc42 protein signal transduction | 2 | 1.78 | 2.11 |
| GO:0030155 | regulation of cell adhesion | 10 | 0.53 | 2.10 |
| GO:0031325 | positive regulation of cellular metabolic process | 27 | 0.27 | 2.10 |
| GO:1902042 | negative regulation of extrinsic apoptotic signaling pathway via death domain receptors | 3 | 1.24 | 2.10 |
| GO:0044092 | negative regulation of molecular function | 14 | 0.42 | 2.08 |
| GO:0019217 | regulation of fatty acid metabolic process | 4 | 0.99 | 2.08 |
| GO:0030100 | regulation of endocytosis | 6 | 0.74 | 2.08 |
| GO:0048523 | negative regulation of cellular process | 35 | 0.22 | 2.07 |
| GO:0002224 | toll-like receptor signaling pathway | 4 | 0.99 | 2.06 |
| GO:0032501 | multicellular organismal process | 46 | 0.17 | 2.06 |
| GO:1903076 | regulation of protein localization to plasma membrane | 4 | 0.99 | 2.06 |
| GO:0034371 | chylomicron remodeling | 2 | 1.72 | 2.04 |
| GO:0050708 | regulation of protein secretion | 8 | 0.6 | 2.04 |
| GO:0071394 | cellular response to testosterone stimulus | 2 | 1.72 | 2.04 |
| GO:0006950 | response to stress | 28 | 0.26 | 2.04 |
| GO:0051240 | positive regulation of multicellular organismal process | 17 | 0.36 | 2.04 |
| GO:0030199 | collagen fibril organization | 3 | 1.21 | 2.03 |
| GO:0007154 | cell communication | 39 | 0.2 | 2.02 |
| GO:0009893 | positive regulation of metabolic process | 28 | 0.25 | 2.02 |
| GO:0051173 | positive regulation of nitrogen compound metabolic process | 26 | 0.27 | 2.02 |
| GO:0048699 | generation of neurons | 16 | 0.37 | 2.01 |
| GO:2001242 | regulation of intrinsic apoptotic signaling pathway | 5 | 0.82 | 2.01 |
| GO:0030154 | cell differentiation | 29 | 0.25 | 2.00 |
| GO:0050714 | positive regulation of protein secretion | 6 | 0.72 | 2.00 |
| GO:0042221 | response to chemical | 33 | 0.22 | 2.00 |
| GO:0090087 | regulation of peptide transport | 10 | 0.51 | 1.99 |
| GO:0016584 | nucleosome positioning | 2 | 1.67 | 1.98 |
| GO:0043589 | skin morphogenesis | 2 | 1.67 | 1.98 |

| Term ID # | Term Description | Count | Enrichment | -Log10PV |
|---|---|---|---|---|
| GO:1905475 | regulation of protein localization to membrane | 5 | 0.81 | 1.97 |
| GO:0051346 | negative regulation of hydrolase activity | 8 | 0.58 | 1.97 |
| GO:0051641 | cellular localization | 21 | 0.31 | 1.97 |
| GO:0010038 | response to metal ion | 7 | 0.64 | 1.96 |
| GO:0051049 | regulation of transport | 18 | 0.34 | 1.96 |
| GO:0120036 | plasma membrane bounded cell projection organization | 13 | 0.42 | 1.96 |
| GO:0031032 | actomyosin structure organization | 4 | 0.94 | 1.94 |
| GO:0031647 | regulation of protein stability | 6 | 0.7 | 1.93 |
| GO:0006959 | humoral immune response | 6 | 0.7 | 1.92 |
| GO:0017157 | regulation of exocytosis | 5 | 0.79 | 1.92 |
| GO:0034378 | chylomicron assembly | 2 | 1.62 | 1.92 |
| GO:0070486 | leukocyte aggregation | 2 | 1.62 | 1.92 |
| GO:0007399 | nervous system development | 21 | 0.3 | 1.92 |
| GO:0045862 | positive regulation of proteolysis | 7 | 0.63 | 1.92 |
| GO:0048585 | negative regulation of response to stimulus | 16 | 0.36 | 1.86 |
| GO:0001568 | blood vessel development | 8 | 0.56 | 1.84 |
| GO:0050790 | regulation of catalytic activity | 21 | 0.29 | 1.83 |
| GO:0002764 | immune response-regulating signaling pathway | 7 | 0.61 | 1.82 |
| GO:0006956 | complement activation | 3 | 1.11 | 1.82 |
| GO:0010896 | regulation of triglyceride catabolic process | 2 | 1.54 | 1.82 |
| GO:0033700 | phospholipid efflux | 2 | 1.54 | 1.82 |
| GO:0043149 | stress fiber assembly | 2 | 1.54 | 1.82 |
| GO:0051222 | positive regulation of protein transport | 7 | 0.61 | 1.82 |
| GO:1903532 | positive regulation of secretion by cell | 7 | 0.61 | 1.82 |
| GO:0010604 | positive regulation of macromolecule metabolic process | 26 | 0.25 | 1.80 |
| GO:0006909 | phagocytosis | 5 | 0.75 | 1.79 |
| GO:0071363 | cellular response to growth factor stimulus | 8 | 0.55 | 1.79 |
| GO:0050994 | regulation of lipid catabolic process | 3 | 1.09 | 1.79 |
| GO:0006957 | complement activation, alternative pathway | 2 | 1.51 | 1.77 |
| GO:0097320 | plasma membrane tubulation | 2 | 1.51 | 1.77 |
| GO:1903729 | regulation of plasma membrane organization | 2 | 1.51 | 1.77 |
| GO:0048731 | system development | 32 | 0.21 | 1.76 |
| GO:0050808 | synapse organization | 5 | 0.75 | 1.76 |
| GO:0071900 | regulation of protein serine/threonine kinase activity | 8 | 0.54 | 1.74 |
| GO:0032271 | regulation of protein polymerization | 5 | 0.74 | 1.74 |
| GO:0010676 | positive regulation of cellular carbohydrate metabolic process | 3 | 1.07 | 1.73 |
| GO:0010035 | response to inorganic substance | 8 | 0.54 | 1.73 |
| GO:0034764 | positive regulation of transmembrane transport | 5 | 0.73 | 1.73 |
| GO:0051046 | regulation of secretion | 10 | 0.46 | 1.73 |
| GO:0070208 | protein heterotrimerization | 2 | 1.48 | 1.73 |
| GO:0070374 | positive regulation of ERK1 and ERK2 cascade | 5 | 0.73 | 1.71 |
| GO:0007169 | transmembrane receptor protein tyrosine kinase signaling pathway | 8 | 0.53 | 1.70 |
| GO:0003012 | muscle system process | 6 | 0.64 | 1.68 |
| GO:0046628 | positive regulation of insulin receptor signaling pathway | 2 | 1.45 | 1.68 |
| GO:0052548 | regulation of endopeptidase activity | 7 | 0.57 | 1.68 |
| GO:0060192 | negative regulation of lipase activity | 2 | 1.45 | 1.68 |
| GO:0071801 | regulation of podosome assembly | 2 | 1.45 | 1.68 |
| GO:0080182 | histone H3-K4 trimethylation | 2 | 1.45 | 1.68 |
| GO:2001028 | positive regulation of endothelial cell chemotaxis | 2 | 1.45 | 1.68 |
| GO:0017144 | drug metabolic process | 9 | 0.48 | 1.68 |
| GO:0048732 | gland development | 7 | 0.57 | 1.68 |
| GO:0051223 | regulation of protein transport | 9 | 0.48 | 1.68 |
| GO:0031333 | negative regulation of protein complex assembly | 4 | 0.84 | 1.68 |

| Term ID # | Term Description | Count | Enrichment | -Log10PV |
|---|---|---|---|---|
| GO:0032272 | negative regulation of protein polymerization | 3 | 1.04 | 1.68 |
| GO:0010243 | response to organonitrogen compound | 11 | 0.42 | 1.67 |
| GO:0000902 | cell morphogenesis | 9 | 0.48 | 1.67 |
| GO:0007051 | spindle organization | 4 | 0.84 | 1.67 |
| GO:0043280 | positive regulation of cysteine-type endopeptidase activity involved in apoptotic process | 4 | 0.84 | 1.67 |
| GO:0048519 | negative regulation of biological process | 36 | 0.18 | 1.67 |
| GO:0099175 | regulation of postsynapse organization | 3 | 1.03 | 1.67 |
| GO:0001956 | positive regulation of neurotransmitter secretion | 2 | 1.42 | 1.66 |
| GO:0032233 | positive regulation of actin filament bundle assembly | 3 | 1.02 | 1.65 |
| GO:0051129 | negative regulation of cellular component organization | 9 | 0.48 | 1.65 |
| GO:0071230 | cellular response to amino acid stimulus | 3 | 1.02 | 1.65 |
| GO:0050806 | positive regulation of synaptic transmission | 4 | 0.82 | 1.65 |
| GO:0009968 | negative regulation of signal transduction | 13 | 0.37 | 1.64 |
| GO:0050789 | regulation of biological process | 66 | 0.1 | 1.63 |
| GO:0034375 | high-density lipoprotein particle remodeling | 2 | 1.39 | 1.63 |
| GO:0043691 | reverse cholesterol transport | 2 | 1.39 | 1.63 |
| GO:0045725 | positive regulation of glycogen biosynthetic process | 2 | 1.39 | 1.63 |
| GO:1903578 | regulation of ATP metabolic process | 3 | 1.01 | 1.63 |
| GO:0060341 | regulation of cellular localization | 10 | 0.44 | 1.62 |
| GO:0035296 | regulation of tube diameter | 4 | 0.81 | 1.62 |
| GO:0051592 | response to calcium ion | 4 | 0.81 | 1.62 |
| GO:0051707 | response to other organism | 13 | 0.37 | 1.61 |
| GO:0120031 | plasma membrane bounded cell projection assembly | 7 | 0.55 | 1.61 |
| GO:0035023 | regulation of Rho protein signal transduction | 4 | 0.81 | 1.60 |
| GO:0001503 | ossification | 5 | 0.69 | 1.60 |
| GO:0042423 | catecholamine biosynthetic process | 2 | 1.37 | 1.60 |
| GO:1903421 | regulation of synaptic vesicle recycling | 2 | 1.37 | 1.60 |
| GO:0031056 | regulation of histone modification | 4 | 0.8 | 1.58 |
| GO:0032717 | negative regulation of interleukin-8 production | 2 | 1.35 | 1.56 |
| GO:1900273 | positive regulation of long-term synaptic potentiation | 2 | 1.35 | 1.56 |
| GO:0010941 | regulation of cell death | 16 | 0.31 | 1.55 |
| GO:0097746 | regulation of blood vessel diameter | 4 | 0.79 | 1.55 |
| GO:0051716 | cellular response to stimulus | 42 | 0.15 | 1.54 |
| GO:0042981 | regulation of apoptotic process | 15 | 0.32 | 1.53 |
| GO:0051004 | regulation of lipoprotein lipase activity | 2 | 1.32 | 1.53 |
| GO:0070584 | mitochondrion morphogenesis | 2 | 1.32 | 1.53 |
| GO:1900371 | regulation of purine nucleotide biosynthetic process | 3 | 0.96 | 1.53 |
| GO:1902176 | negative regulation of oxidative stress-induced intrinsic apoptotic signaling pathway | 2 | 1.32 | 1.53 |
| GO:1903530 | regulation of secretion by cell | 9 | 0.45 | 1.53 |
| GO:0048856 | anatomical structure development | 36 | 0.17 | 1.53 |
| GO:0030833 | regulation of actin filament polymerization | 4 | 0.78 | 1.52 |
| GO:0051225 | spindle assembly | 3 | 0.96 | 1.52 |
| GO:0034394 | protein localization to cell surface | 2 | 1.3 | 1.51 |
| GO:0072359 | circulatory system development | 10 | 0.42 | 1.51 |
| GO:0000122 | negative regulation of transcription by RNA polymerase II | 10 | 0.42 | 1.50 |
| GO:0002757 | immune response-activating signal transduction | 6 | 0.58 | 1.50 |
| GO:0022898 | regulation of transmembrane transporter activity | 5 | 0.66 | 1.50 |
| GO:0031324 | negative regulation of cellular metabolic process | 21 | 0.25 | 1.50 |
| GO:0043086 | negative regulation of catalytic activity | 10 | 0.42 | 1.50 |
| GO:0062012 | regulation of small molecule metabolic process | 6 | 0.58 | 1.50 |
| GO:0032535 | regulation of cellular component size | 6 | 0.58 | 1.50 |
| GO:0003008 | system process | 17 | 0.29 | 1.50 |
| GO:0051172 | negative regulation of nitrogen compound metabolic process | 20 | 0.26 | 1.49 |

| Term ID # | Term Description | Count | Enrichment | -Log10PV |
|---|---|---|---|---|
| GO:0002526 | acute inflammatory response | 3 | 0.94 | 1.49 |
| GO:0050995 | negative regulation of lipid catabolic process | 2 | 1.28 | 1.49 |
| GO:0006952 | defense response | 13 | 0.35 | 1.48 |
| GO:0071345 | cellular response to cytokine stimulus | 11 | 0.39 | 1.48 |
| GO:0016310 | phosphorylation | 13 | 0.34 | 1.48 |
| GO:0051241 | negative regulation of multicellular organismal process | 12 | 0.36 | 1.47 |
| GO:0071902 | positive regulation of protein serine/threonine kinase activity | 6 | 0.57 | 1.47 |
| GO:0030325 | adrenal gland development | 2 | 1.26 | 1.46 |
| GO:0090140 | regulation of mitochondrial fission | 2 | 1.26 | 1.46 |
| GO:0030705 | cytoskeleton-dependent intracellular transport | 4 | 0.75 | 1.45 |
| GO:0010951 | negative regulation of endopeptidase activity | 5 | 0.64 | 1.45 |
| GO:0033344 | cholesterol efflux | 2 | 1.24 | 1.43 |
| GO:0051194 | positive regulation of cofactor metabolic process | 2 | 1.24 | 1.43 |
| GO:0045861 | negative regulation of proteolysis | 6 | 0.56 | 1.42 |
| GO:0009894 | regulation of catabolic process | 10 | 0.4 | 1.42 |
| GO:0007265 | Ras protein signal transduction | 4 | 0.73 | 1.42 |
| GO:1903827 | regulation of cellular protein localization | 7 | 0.5 | 1.41 |
| GO:0019433 | triglyceride catabolic process | 2 | 1.23 | 1.40 |
| GO:0050794 | regulation of cellular process | 62 | 0.09 | 1.40 |
| GO:0030838 | positive regulation of actin filament polymerization | 3 | 0.89 | 1.40 |
| GO:0048259 | regulation of receptor-mediated endocytosis | 3 | 0.89 | 1.40 |
| GO:0030307 | positive regulation of cell growth | 4 | 0.72 | 1.39 |
| GO:0032502 | developmental process | 37 | 0.16 | 1.39 |
| GO:0045927 | positive regulation of growth | 5 | 0.62 | 1.39 |
| GO:1901700 | response to oxygen-containing compound | 14 | 0.31 | 1.39 |
| GO:0030041 | actin filament polymerization | 2 | 1.21 | 1.39 |
| GO:0043534 | blood vessel endothelial cell migration | 2 | 1.21 | 1.39 |
| GO:0051220 | cytoplasmic sequestering of protein | 2 | 1.21 | 1.39 |
| GO:0032990 | cell part morphogenesis | 7 | 0.49 | 1.38 |
| GO:0050807 | regulation of synapse organization | 4 | 0.72 | 1.38 |
| GO:0001649 | osteoblast differentiation | 3 | 0.88 | 1.38 |
| GO:0007568 | aging | 5 | 0.62 | 1.38 |
| GO:0043066 | negative regulation of apoptotic process | 10 | 0.39 | 1.38 |
| GO:0043255 | regulation of carbohydrate biosynthetic process | 3 | 0.88 | 1.38 |
| GO:0045088 | regulation of innate immune response | 6 | 0.54 | 1.38 |
| GO:0002697 | regulation of immune effector process | 6 | 0.54 | 1.38 |
| GO:0007163 | establishment or maintenance of cell polarity | 4 | 0.71 | 1.38 |
| GO:0031349 | positive regulation of defense response | 6 | 0.54 | 1.37 |
| GO:0048167 | regulation of synaptic plasticity | 4 | 0.71 | 1.37 |
| GO:0051354 | negative regulation of oxidoreductase activity | 2 | 1.19 | 1.37 |
| GO:2001171 | positive regulation of ATP biosynthetic process | 2 | 1.19 | 1.37 |
| GO:0006638 | neutral lipid metabolic process | 3 | 0.87 | 1.36 |
| GO:0050821 | protein stabilization | 4 | 0.7 | 1.36 |
| GO:0071417 | cellular response to organonitrogen compound | 7 | 0.48 | 1.36 |
| GO:0090162 | establishment of epithelial cell polarity | 2 | 1.18 | 1.35 |
| GO:0002758 | innate immune response-activating signal transduction | 4 | 0.7 | 1.35 |
| GO:0031329 | regulation of cellular catabolic process | 9 | 0.41 | 1.34 |
| GO:0007179 | transforming growth factor beta receptor signaling pathway | 3 | 0.86 | 1.34 |
| GO:0008015 | blood circulation | 6 | 0.53 | 1.34 |
| GO:0002684 | positive regulation of immune system process | 10 | 0.38 | 1.33 |
| GO:0009892 | negative regulation of metabolic process | 22 | 0.22 | 1.33 |
| GO:0045892 | negative regulation of transcription, DNA-templated | 12 | 0.33 | 1.33 |
| GO:0060191 | regulation of lipase activity | 3 | 0.85 | 1.32 |
| GO:0009888 | tissue development | 15 | 0.29 | 1.32 |

| Term ID # | Term Description | Count | Enrichment | -Log10PV |
|---|---|---|---|---|
| GO:0045922 | negative regulation of fatty acid metabolic process | 2 | 1.15 | 1.31 |
| GO:0044089 | positive regulation of cellular component biogenesis | 7 | 0.47 | 1.31 |
| GO:0046890 | regulation of lipid biosynthetic process | 4 | 0.68 | 1.31 |
| GO:0031327 | negative regulation of cellular biosynthetic process | 14 | 0.3 | 1.31 |
| GO:0048666 | neuron development | 9 | 0.4 | 1.30 |
| GO:0045055 | regulated exocytosis | 25 | 0.88 | 11.30 |
| **Decreased in post LT fibrosis** | | | | |
| GO:0002576 | platelet degranulation | 10 | 1.49 | 8.67 |
| GO:0007010 | cytoskeleton organization | 16 | 0.83 | 6.24 |
| GO:0016192 | vesicle-mediated transport | 20 | 0.67 | 6.14 |
| GO:0045055 | regulated exocytosis | 13 | 0.88 | 5.38 |
| GO:0030029 | actin filament-based process | 10 | 0.91 | 4.05 |
| GO:0030036 | actin cytoskeleton organization | 9 | 0.93 | 3.77 |
| GO:0016043 | cellular component organization | 29 | 0.35 | 3.66 |
| GO:0022607 | cellular component assembly | 19 | 0.51 | 3.62 |
| GO:0007229 | integrin-mediated signaling pathway | 5 | 1.38 | 3.48 |
| GO:0097435 | supramolecular fiber organization | 8 | 0.92 | 3.28 |
| GO:0006953 | acute-phase response | 4 | 1.55 | 3.22 |
| GO:0006810 | transport | 24 | 0.37 | 2.92 |
| GO:0035296 | regulation of tube diameter | 5 | 1.19 | 2.80 |
| GO:0090066 | regulation of anatomical structure size | 8 | 0.84 | 2.80 |
| GO:0042730 | fibrinolysis | 3 | 1.76 | 2.74 |
| GO:0051179 | localization | 27 | 0.31 | 2.74 |
| GO:0097746 | regulation of blood vessel diameter | 5 | 1.16 | 2.74 |
| GO:0030193 | regulation of blood coagulation | 4 | 1.32 | 2.57 |
| GO:0031589 | cell-substrate adhesion | 5 | 1.09 | 2.52 |
| GO:0003012 | muscle system process | 6 | 0.92 | 2.37 |
| GO:0006996 | organelle organization | 19 | 0.38 | 2.31 |
| GO:0002376 | immune system process | 16 | 0.43 | 2.21 |
| GO:0007015 | actin filament organization | 5 | 1 | 2.19 |
| GO:2001237 | negative regulation of extrinsic apoptotic signaling pathway | 4 | 1.19 | 2.19 |
| GO:0007044 | cell-substrate junction assembly | 3 | 1.48 | 2.17 |
| GO:0043152 | induction of bacterial agglutination | 2 | 2.12 | 2.17 |
| GO:0001775 | cell activation | 10 | 0.59 | 2.12 |
| GO:0006952 | defense response | 11 | 0.55 | 2.12 |
| GO:0007160 | cell-matrix adhesion | 4 | 1.13 | 2.05 |
| GO:0006897 | endocytosis | 7 | 0.74 | 2.01 |
| GO:0065003 | protein-containing complex assembly | 12 | 0.5 | 2.01 |
| GO:0043933 | protein-containing complex subunit organization | 13 | 0.47 | 2.00 |
| GO:0007167 | enzyme linked receptor protein signaling pathway | 8 | 0.66 | 2.00 |
| GO:0008015 | blood circulation | 6 | 0.81 | 2.00 |
| GO:0033622 | integrin activation | 2 | 1.95 | 2.00 |
| GO:0006936 | muscle contraction | 5 | 0.91 | 1.97 |
| GO:0034329 | cell junction assembly | 4 | 1.07 | 1.97 |
| GO:0031532 | actin cytoskeleton reorganization | 3 | 1.32 | 1.92 |
| GO:0031639 | plasminogen activation | 2 | 1.86 | 1.91 |
| GO:0051640 | organelle localization | 7 | 0.69 | 1.84 |
| GO:0034116 | positive regulation of heterotypic cell-cell adhesion | 2 | 1.79 | 1.81 |
| GO:0051641 | cellular localization | 14 | 0.41 | 1.78 |
| GO:0140056 | organelle localization by membrane tethering | 4 | 1 | 1.77 |
| GO:0007017 | microtubule-based process | 7 | 0.66 | 1.76 |
| GO:0098657 | import into cell | 7 | 0.66 | 1.75 |
| GO:1903524 | positive regulation of blood circulation | 3 | 1.23 | 1.75 |
| GO:0030198 | extracellular matrix organization | 5 | 0.83 | 1.71 |
| GO:0048268 | clathrin coat assembly | 2 | 1.67 | 1.67 |
| GO:0001568 | blood vessel development | 6 | 0.71 | 1.65 |
| GO:0008217 | regulation of blood pressure | 4 | 0.96 | 1.65 |

| Term ID # | Term Description | Count | Enrichment | -Log10PV |
|---|---|---|---|---|
| GO:0002064 | epithelial cell development | 4 | 0.95 | 1.65 |
| GO:0050878 | regulation of body fluid levels | 6 | 0.7 | 1.60 |
| GO:0051258 | protein polymerization | 3 | 1.16 | 1.60 |
| GO:0045087 | innate immune response | 7 | 0.62 | 1.56 |
| GO:0003008 | system process | 12 | 0.42 | 1.56 |
| GO:0032970 | regulation of actin filament-based process | 5 | 0.77 | 1.55 |
| GO:0045185 | maintenance of protein location | 3 | 1.1 | 1.47 |
| GO:0031032 | actomyosin structure organization | 3 | 1.09 | 1.47 |
| GO:0070527 | platelet aggregation | 2 | 1.5 | 1.46 |
| GO:0022407 | regulation of cell-cell adhesion | 5 | 0.74 | 1.43 |
| GO:0030194 | positive regulation of blood coagulation | 2 | 1.49 | 1.43 |
| GO:0032967 | positive regulation of collagen biosynthetic process | 2 | 1.49 | 1.43 |
| GO:0007166 | cell surface receptor signaling pathway | 13 | 0.37 | 1.43 |
| GO:0072378 | blood coagulation, fibrin clot formation | 2 | 1.47 | 1.43 |
| GO:2000352 | negative regulation of endothelial cell apoptotic process | 2 | 1.46 | 1.42 |
| GO:0045595 | regulation of cell differentiation | 11 | 0.41 | 1.39 |
| GO:0030866 | cortical actin cytoskeleton organization | 2 | 1.43 | 1.37 |
| GO:0110053 | regulation of actin filament organization | 4 | 0.83 | 1.36 |
| GO:0006911 | phagocytosis, engulfment | 2 | 1.41 | 1.35 |
| GO:1900026 | positive regulation of substrate adhesion-dependent cell spreading | 2 | 1.4 | 1.33 |
| GO:0045907 | positive regulation of vasoconstriction | 2 | 1.38 | 1.31 |

**Supplemental Table 6. Feeding, growth and lipid accumulation caused by HFD in C57Bl6/J and AJ mice.**

| | *LFD* | | *HFD* | |
|---|---|---|---|---|
| | C57Bl6/J | AJ | C57Bl6/J | AJ |
| *Food consumption (g/d)* | 2.7±0.1 | 2.6±0.1 | 2.7±0.1 | 2.5±0.2 |
| *BW gain (g/wk)* | 0.4±0.1 | 0.6±0.1 | 1.5±0.1[a] | 0.6±0.1[b] |
| *LW (% of BW)* | 3.5±0.1 | 4.1±0.1 | 6.5±0.4[a] | 4.6±0.1[b] |
| *Hepatic TG (µg/mg liver)* | 5.3±2.1 | 5.4±1.3 | 140±41[a] | 17.6±5.5[b] |
| *Hepatic FFA (ng/mg liver)* | 0.1±0.1 | 0.2±0.1 | 7.5±1.9[a] | 1.7±1.0[b] |
| *Hepatic TC (µg/mg liver)* | 0.4±0.1 | 0.9±0.1 | 7.0±1.6[a] | 5.4±1.5[a] |

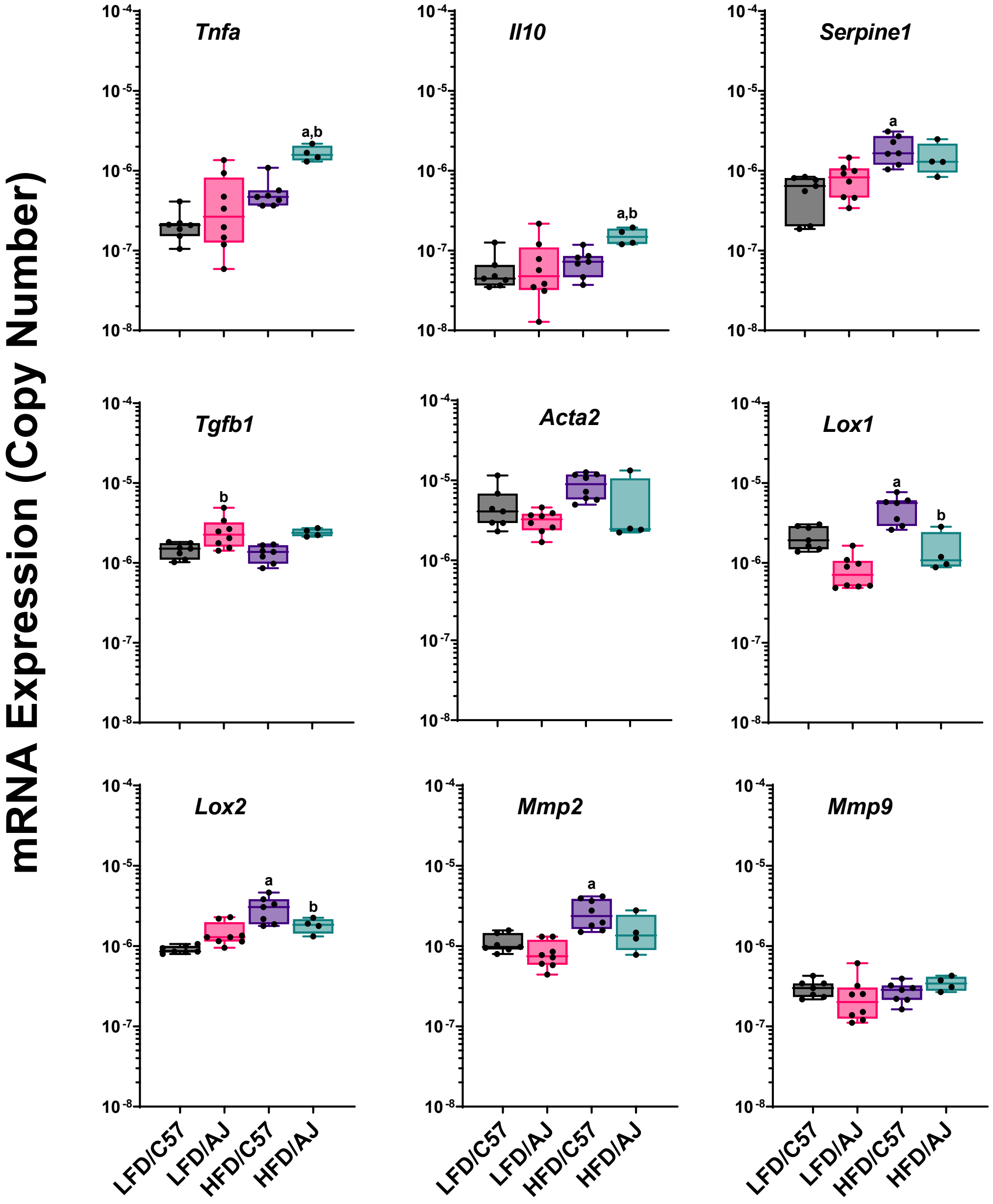


Supplemental Figure 1-Li et al.
